# RatA is not a toxin but serves as a ubiquinone shuttle

**DOI:** 10.64898/2026.05.04.722385

**Authors:** Michel Fasnacht, Leila Jensen, Denise Schratt, Isabella Moll

**Affiliations:** Max Perutz Labs, Vienna BioCenter, 1030 Vienna, Austria; University of Vienna, Vienna, Austria; University of Vienna, Doctoral School in Microbiology and Environmental Science, Vienna, Austria

## Abstract

Conflicting roles have been proposed for the *E. coli* protein RatA. Initially described as a ribosome targeting toxin, a later report pronounced it the bacterial homologue to the inner mitochondrial membrane protein Coq10. Coq10 proteins are conserved from prokaryotes to human and implicated to serve a lipid chaperone role in the biosynthesis of ubiquinone, a crucial electron carrier during aerobic respiration. We recently identified that the contradictory results published for RatA can be attributed to a mis-annotation of the gene in the reference genome. Here, we further elucidate the molecular function of RatA. We clarify that RatA is not a toxin but serves as a lipid shuttle for ubiquinone from its cytosolic biosynthesis complex to the inner membrane. Furthermore, we show that the loss of RatA results in an impaired, but not abolished electron transport chain and demonstrate broad metabolic adaptations of the cells as a consequence. Therefore, we propose to rename RatA to UbiM to reflect its function and to be in accordance with the naming convention of other ubiquinone biosynthesis proteins.

## Introduction

After being first proposed in 1961^1^, the chemiosmotic theory of transmembrane differences in proton concentrations driving the synthesis of ATP has become well established textbook knowledge. Formation of the required proton gradient is powered by the flow of electrons through the electron transport chain (ETC), a series of membrane-bound protein complexes and electron carriers. In *E. coli*, the ETC is located in the plasma membrane, whereas in eukaryotes it is located in the inner mitochondrial membrane. In both cases, during aerobic growth, ubiquinone (also known as coenzyme Q, CoQ, or simply Q) serves as an electron shuttle between different respiratory complexes (Figure 1A). Ubiquinone contains a redox-active benzoquinone head group and a polyisoprenoid tail that differs in length between organisms (Figure 1B). In *E. coli*, the tail consists of eight isoprenoid units (Q_8_). Ubiquinone biosynthesis has been studied intensively, and responsible enzymes are often functionally conserved even among distantly related organisms, yet many open questions remain^2–6^. In mitochondria, the synthesis is performed by a series of enzymes in the still ill-defined complex Q (or CoQ-synthome)^3,4,7–9^. In *E. coli*, Q_8_ synthesis is catalyzed by the enzymes encoded in the *ubi* genes^2^. In a recent study, Chehade et al. further characterized the synthetic complexes formed by the Ubi proteins^10^. Q_8_ synthesis begins in the plasma membrane, but, rather unexpectedly based on the strong hydrophobic characteristics of ubiquinone, ends in the cytosol. In fact, the majority of the head-modifying reactions are catalyzed in the cytosol by an approximately 1 MDa complex consisting of 5 different Ubi proteins interacting with an UbiJ-UbiK scaffold^10^. Importantly, the exact stoichiometry of the complex as well as the mechanistic details of the delivery of the ubiquinone precursor molecules to the metabolon and of the transport of Q_8_ back to the plasma membrane remain elusive (Figure 1A).

**Figure 1.**
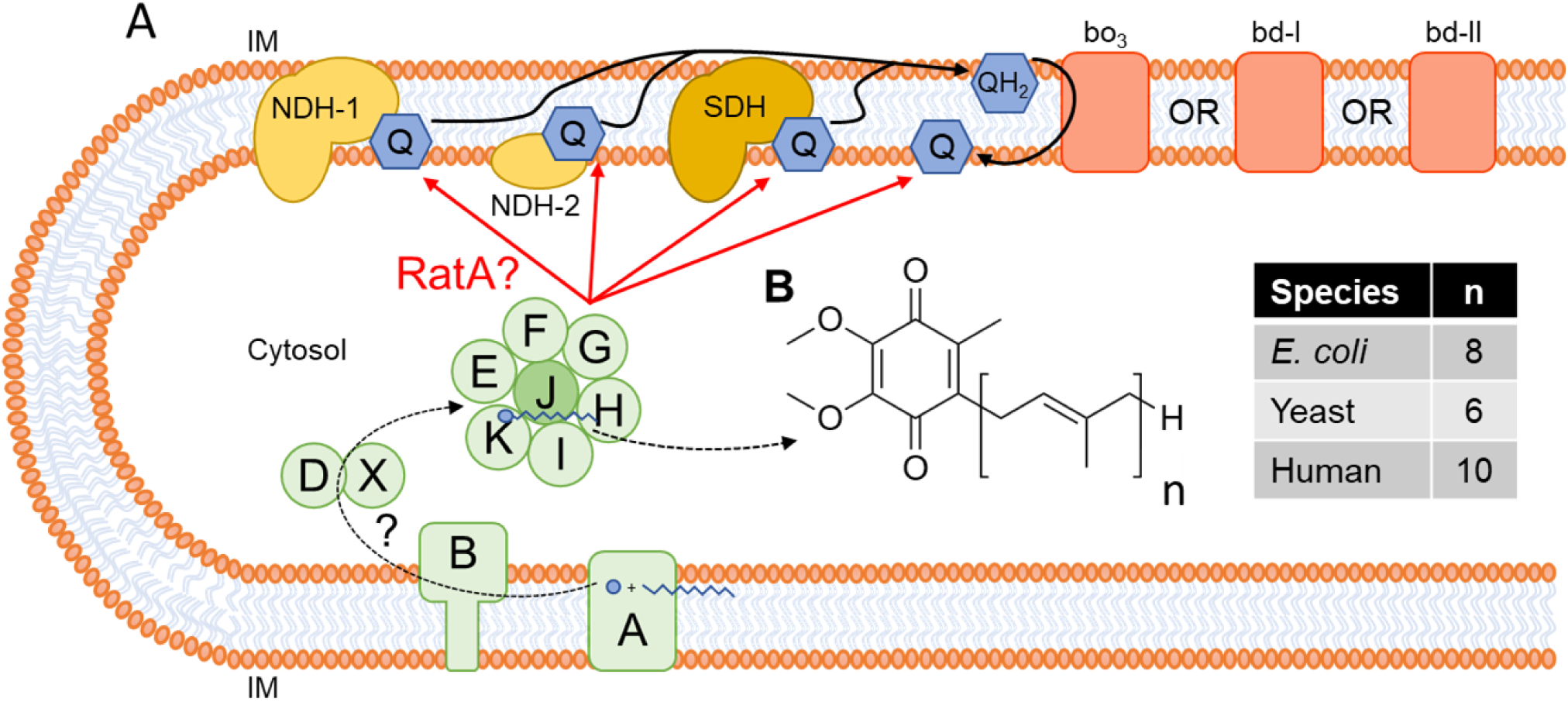
Simplified model of the synthesis and function of ubiquinone in *E. coli*. **A** During aerobic respiration, ubiquinone (Q) and ubiquinol (QH_2_) serve as an electron shuttle between the initial oxidoreductases (NDH-1: NADH:quinone oxidoreductase I, NDH-2: NADH:quinone oxidoreductase II, SDH: succinate:quinone oxidoreductase) and the terminal oxidoreductases (cytochrome bo_3_, bd-I, or bd-II). In *E. coli*, the synthesis of ubiquinone begins in the inner membrane (IM) and is finalized in an approximately 1 MDa cytosolic complex consisting of 7 different Ubi proteins (green circles) with a so far uncharacterized stoichiometry. Currently, it has not been determined how precursor molecules are delivered from the membrane to the metabolon and how the finished ubiquinone is transported from the complex to the inner membrane. **B** The chemical structure of ubiquinone varies between species in the respective length of the isoprenoid tail.

The ribosome associated toxin A (RatA) of the *E. coli* K-12 strains was initially described as the toxin moiety of a toxin-antitoxin (TA) pair that targets the 50S ribosomal subunit to inhibit subunits association and thereby blocks the initiation step of translation^11,12^. However, later studies on its homologue PasT yielded conflicting results. PasT is found in uropathogenic *E. coli* (UPEC) strains and differs from RatA in only two amino acids (N90S and E111D). While an initial study further confirmed a toxic effect upon overexpression of *pasT* and described a plethora of phenotypes related to stress resistance and infectivity in a *pasT* deletion strain^13^, a later study strongly suggested that PasT is the bacterial homologue of the mitochondrial protein Coq10^14^. Coq10-related proteins are found from prokaryotes to humans^15,16^ (Supplementary Figure 1B-E), but are best studied in yeast. Coq10 is an inner mitochondrial membrane protein^15,17^, which distributes uniformly among mitochondria^8^. It contains a steroidogenic acute regulatory protein-related lipid transfer (START) domain and can directly bind Q_6_ and several analogs thereof^15,17–19^. While two studies identified impaired *de novo* biosynthesis of Q_6_ specifically in early log phase upon knockout of the gene^16,20^, Coq10 is not essential for ubiquinone synthesis, manifested by *coq10* null mutants generally harboring the same total amount of Q_6_ as the wildtype strains^15,17^. Nevertheless, cells lacking Coq10 still display several phenotypes related to impaired respiratory electron transport^15,17–19^, which can be complemented by Coq10 homologues originating from different organisms^15–17,20^. It is therefore thought that Coq10 serves a conserved lipid chaperone function, potentially delivering newly synthesized ubiquinone to the ETC and/or ensuring the correct positioning of ubiquinone molecules in the ETC. However, direct evidence for the precise role of Coq10 remains missing.

Like in *coq10* null mutants, deletion of *pasT* or *ratA* resulted in an impaired electron transport through the ETC, revealed by a decrease in membrane potential^14^. Furthermore, total Q_8_ levels were not significantly different compared to the parental strains, though a decrease in ubiquinone precursors during exponential growth was observed in UPEC cells devoid of PasT^14^. Lastly, a hypersensitivity to oxidative stress was demonstrated^14^. Importantly, all phenotypes could at least partially be rescued by human Coq10A, providing the strongest evidence that PasT/RatA is the *E. coli* homologue of Coq10^14^. It has been argued that the observed phenotypes upon loss of PasT are likely indirect and downstream effects originating from a broad distortion of bacterial physiology caused by the defective respiration apparatus^21^. Indeed, transcriptome analysis of an avian pathogenic *E. coli* (APEC) strain provided first indications that a multitude of genes belonging to a variety of metabolic pathways were differentially expressed upon deletion of *ratA*^22^. However, the precise function of RatA in the ubiquinone biosynthesis remains to be established

We recently identified that the contradictory results published for RatA being either part of a TA system or the bacterial homologue of Coq10 can be attributed to a mis-annotation of the gene in the reference genome, a phenomenon that could be more widespread for bacterial genes^23^. In this study, we further elucidate the molecular function of RatA. We demonstrate that RatA is not a toxin and experimentally confirm for the first time Q_8_ binding by RatA. We further illustrate that two additional hallmark phenotypes associated with the eukaryotic *coq10* mutant strains are equally found in bacterial *ratA* deletion strains and demonstrate broad metabolic adaptations of *E. coli* cells upon loss of RatA by analyzing both the metabolome and proteome of a *ratA* deletion strain. In contrast to Coq10, we show that RatA is not only present in the inner membrane, but equally resides in the cytosol. Since we also demonstrate interactions of RatA with the ubiquinone synthesis complex, we propose that the functional role of RatA is that of a lipid shuttle transporting mature Q_8_ from the metabolon in the cytosol to the inner membrane and should thus be renamed.

## Results

### RatA_Δ14_ is the only variant synthesized under all conditions tested

Previously, we showed that the reported toxicity of RatA is likely an artifact caused by the incorrect annotation of the start codon of *ratA*^23^. Expression of the annotated sequence of *ratA* yields two RatA variants of differing lengths, the toxic full-length RatA (RatA_fl)_ with an N-terminal extension of 14 amino acids and the shorter, non-toxic RatA_Δ14_^23^. Our data revealed that under optimal, aerobic conditions, expression of *ratA* from its endogenous promoter produces RatA_Δ14_ only^23^. However, it was tempting to speculate that the full-length RatA variant is engendered conditionally governed by an alternative upstream promoter. Hence, we checked for the presence of RatA_fl_ under stress conditions that resulted in significant phenotypes upon *ratA* or *pasT* deletion. More specifically, we included nutrient depletion^13^, oxidative stress^13,21^, and antibiotic treatments^13,21^. As shown in Figure 2, no RatA_fl_ was detected under any of the tested conditions. Fittingly, most of the phenotypes driven by *pasTI* deletion in CFT073 were alleviated with the short variant of PasT originating from methionine 14 (Met14). In contrast, rescuing persister cells formation in the presence of ciprofloxacin required leaky expression of the full-length *pasT* gene^13^. To exclude the possibility of strain-specific effects, we re-examined the response to ciprofloxacin treatment in a CFT073Δ*pasTI* background. Since no sequence differences exist in the region upstream of *pasT* and *ratA*, we likewise employed plasmid pZS*2-ratA-His, but still detected no RatA_fl_ after addition of ciprofloxacin (Supplementary Figure 2). Taken together, our results strongly support the notion that RatA_Δ14_ is the only physiologically relevant RatA variant and will thus refer to RatA_Δ14_ as RatA throughout the remainder of the manuscript unless specified otherwise.

**Figure 2.**
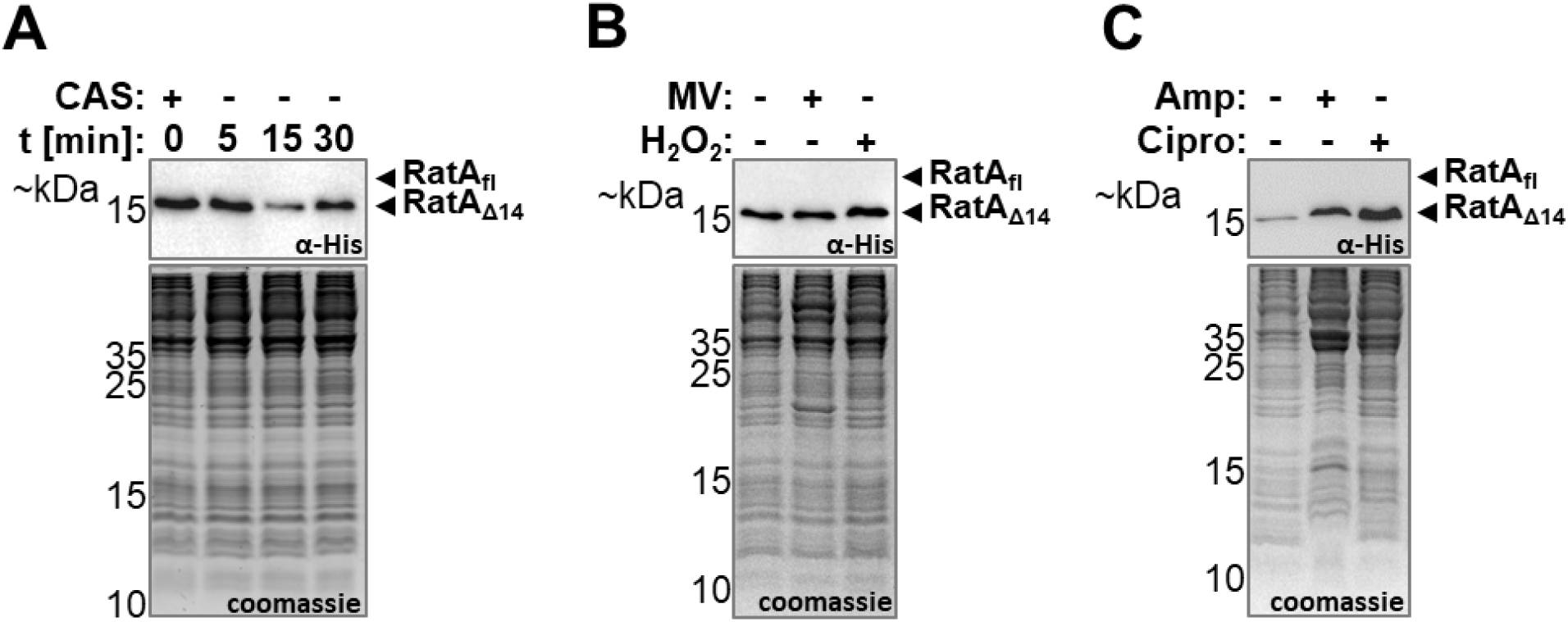
Only the non-toxic RatA_Δ14_ variant is detected regardless of the stress condition. Western blot analysis of samples harvested from the BW25113(Δ*ratA*) pZS*2–ratA–His strain ectopically expressing the wildtype *ratA* gene with a 3′-terminal His-tag sequence under endogenous promoter control to investigate whether the toxic RatA_fl_ variant is detected under specific stress conditions. Coomassie gels served as a loading control. **A** Nutritional stress was induced by the removal of casamino acids (CAS). **B** Oxidative stress was induced either by the addition of 1 mM paraquat (methyl viologen, MV) or 10 mM hydrogen peroxide (H_2_O_2_). **C** Antibiotic stress was induced by the addition of either f.c. 100 µg/ml ampicillin (Amp) or 10 µg/ml ciprofloxacin (Cipro).

### RatA binds ubiquinone in *E. coli*

Structure prediction by AlphaFold^24^ for RatA results in a START domain family protein comprising numerous conserved residues and structural similarities within the hydrophobic pocket to its homologs in *Caulobacter crescentus*, *Saccharomyces cerevisiae*, and humans (Supplementary Figure 1). To experimentally validate binding of Q_8_, we purified RatA via a C-terminal Strep-tag and analyzed the co-purified hydrophobic ligands. As a control, we simultaneously purified Strep-tagged H-NS, a well-studied DNA binding protein of comparable size to RatA without any known lipid binding activity^25^. For normalization, a soluble ubiquinone variant with a shortened isoprenoid tail (Q_2_) was spiked-in to the elution buffer. As shown in Figure 3, we observed a clear enrichment of Q_8_ in the elution fractions containing RatA compared to the H-NS elution fractions.

**Figure 3.**
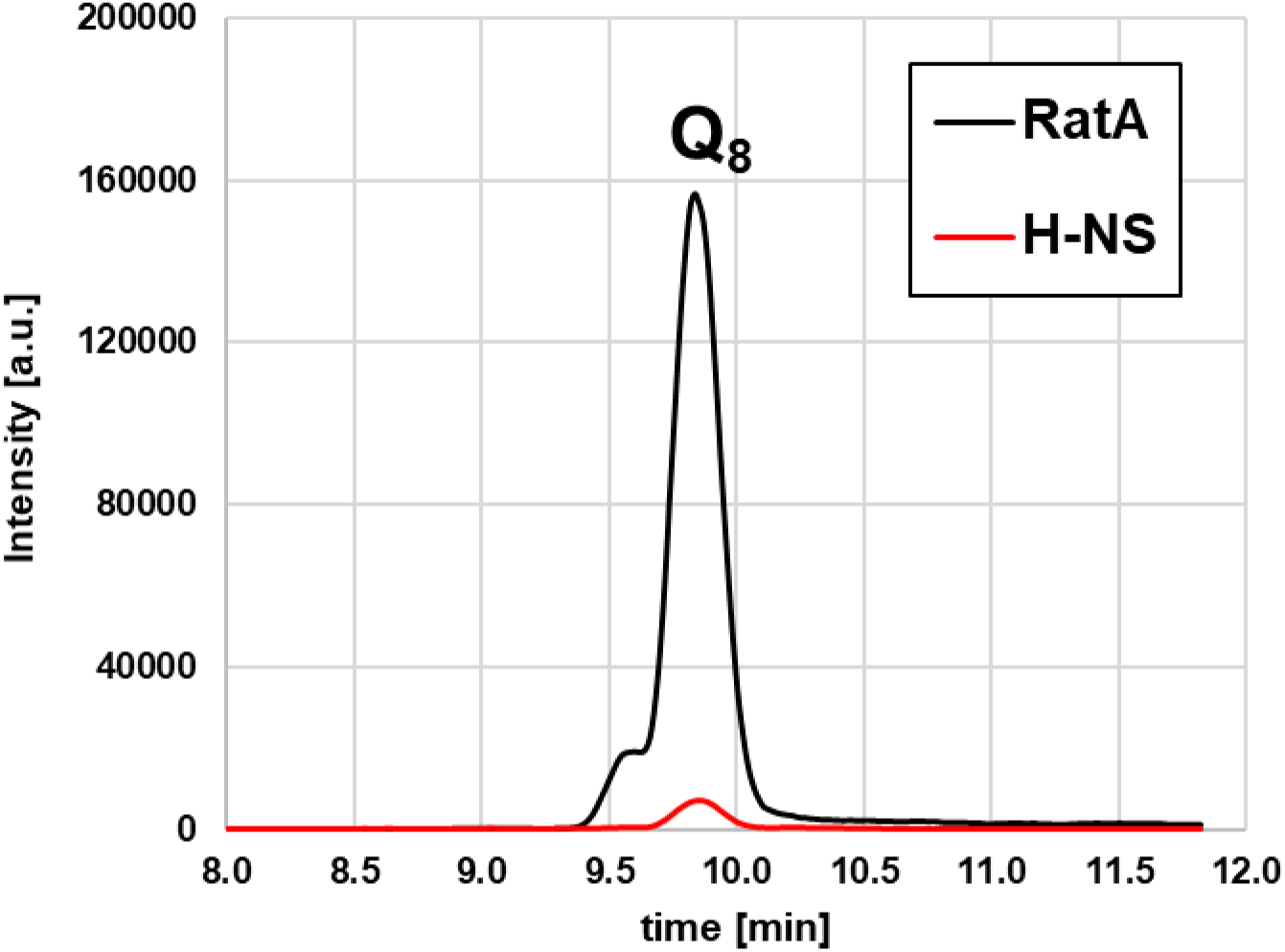
RatA binds Q_8_. Q_8_ binding to RatA was confirmed by extracting the protein via a C-terminal Strep-tag followed by chloroform/methanol extraction of the bound ligands. Q_8_ was detected, confirmed and quantified by LC-MS/MS, employing the selected reaction monitoring (SRM) mode of the mass spectrometer. Shown is a representative elution profile of the liquid chromatograph with Q_8_ eluting after approximately 9.9 minutes. As a control, the DNA-binding protein H-NS was employed. Its peak was normalized to the Q_2_ peaks (data not shown), which was added as a spike-in control to the protein elution buffer.

To identify amino acids critical for Q_8_ binding, we introduced targeted mutations at residues previously implicated in ubiquinone binding^15,19,22^. Two mutations, F43A and W103A, appeared to completely impede RatA functionality, as demonstrated by the small colony size observed for the *ratA* deletion strain (Supplementary Figure 3A, B). Two additional mutations, P45A and V64A, demonstrated a less pronounced reduction in colony size (Supplementary Figure 3A, B), rendering them potentially interesting targets for further investigations. However, all mutations drastically affected protein abundance as determined by western blot analyses (Supplementary Figure 3C, D). Thus, the observed small colony phenotype can be attributed to the reduction in protein abundance rather than to a reduction in Q_8_ binding.

### Deletion of *ratA* in *E. coli* mirrors hallmark phenotypes of *coq10* deletion strains

Comparable to eukaryotic cells lacking Coq10, the *pasT/ratA* deletion mutants of *E. coli* strains CFT073 and MG1655 respectively displayed phenotypes related to impaired respiratory electron transport though with some strain-specific differences. Hence, we first re-examined the effects of *ratA* deletion in our model organism, the *E. coli* K-12 strain BW25113. As evidenced in Figure 4A, the *ratA* deletion strain and its isogenic wildtype strain similarly re-initiate growth in LB medium under optimal, aerobic growth conditions in flasks. However, the deletion strain exhibits a clear growth defect in the early exponential phase, which is partially rescued by the presence of C-terminal His-tagged RatA. This result is in contrast to a previous study, where deletions of *pasT* or *ratA* did not affect growth in LB medium when incubated in an automated plate reader system^13^, but consistent with the growth phenotype observed for the *ratA* deletion in the APEC strain cultivated in flasks^22^. As the surface area to volume ratio differs significantly between flasks and microtiter plates, thereby affecting the aeration of the culture, we compared growth in the plate reader in different volumes of nutrient rich LB medium. As shown in Supplementary Figure 4, the wildtype and the *ratA* deletion strain exhibit a clear growth difference at the beginning of the exponential phase only when incubated in smaller culture volumes. As in flasks, this phenotype can be complemented by ectopic expression of *ratA* under endogenous promoter control (Supplementary Figure 5). Contrary to the previously published results^13^, we observed the same growth phenotype in LB medium upon deletion of the *pasT* gene from the genome of the pathogenic CFT073 *E. coli* strain (Figure 4B).

**Figure 4.**
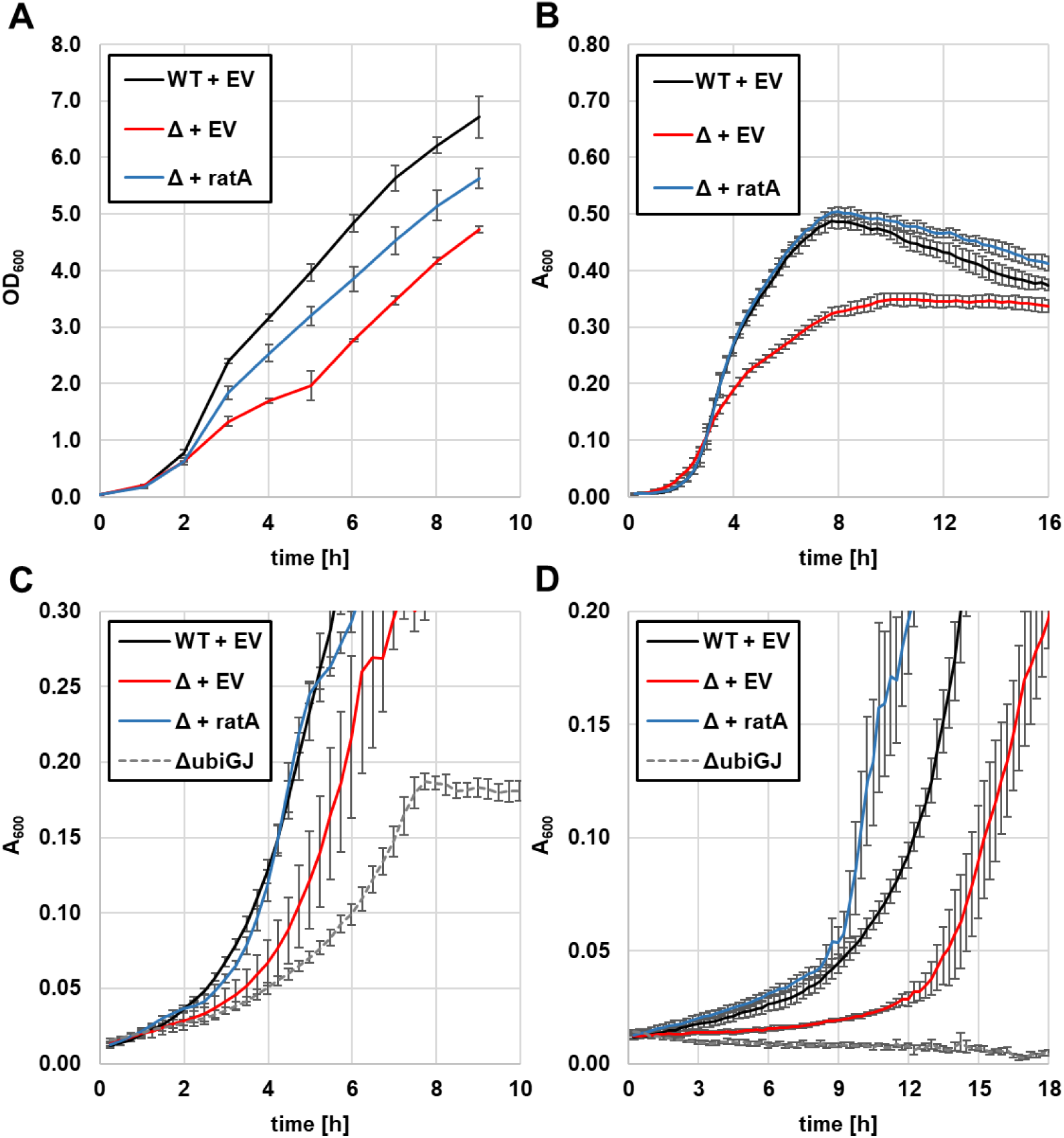
Cells lacking RatA display a growth phenotype that is more pronounced when grown on succinate. **A** The *ratA* deletion strain transformed with the empty vector pZS*2 (Δ + EV, red line) displayed a clear growth phenotype compared to the corresponding wildtype *E. coli* strain BW25113 transformed with the same empty vector (WT + EV, black line) when grown in LB medium in flasks at 37°C while continuously shaking. Ectopically expressing the *ratA* gene in the deletion strain under endogenous promoter control from the pZS*2-ratA-His plasmid (Δ + ratA, blue line) partially restored the observed phenotype. **B** The same growth phenotype compared to its corresponding wildtype was observed upon deletion of the *pasT* gene in the pathogenic *E. coli* strain CFT073 when grown in LB medium in a 96 well plate at 37°C while continuously shaking. Again, introduction of the *ratA* gene by transformation of the pZS*2-ratA-His plasmid into the *pasT* deletion strain restored wildtype growth. **C** Same strains as in A, but grown in 1x M9 + 0.2% glucose in a 96 well plate at 37°C while continuously shaking. As an additional control, a double knockout strain lacking both the *ubiG* and the *ubiJ* gene transformed with the empty vector pZS*2 was employed (ΔubiGJ, grey dotted line) **D** Same strains as in C, but grown in 1x M9 + 0.2% succinate in a 96 well plate at 37°C while continuously shaking. All panels display the average of 3 biological replicates with error bars representing the standard deviation.

LB medium is abundant in a variety of catabolyzable amino acids and a limited amount of fermentable sugars^26^. We thus hypothesized that if the deletion of *ratA* negatively affects the ETC, the deletion strain should exhibit an even stronger growth defect if grown on a non-fermentable sugar as the sole carbon source. We therefore monitored growth in M9 minimal medium supplemented with either glucose or succinate. As control, we included a *ubiG/ubiJ* double deletion strain, which is deficient in ubiquinone synthesis and should thus not grow on non-fermentable sugars. In the presence of glucose, strains exhibit similar growth patterns as in the LB medium (Figure 4C). Compared to the wildtype strain, a clear growth phenotype of cells lacking RatA was observed after the lag phase, while the *ubiG/ubiJ* double deletion strain was even more impaired in growth (Figure 4C). As expected, in the presence of the TCA cycle intermediate succinate, we observed no growth of the *ubiG/ubiJ* double deletion mutant. In contrast, the *ratA* deficient cells displayed a significantly longer lag phase compared to the wildtype cells, but did ultimately manage to grow on succinate (Figure 4D). Interestingly, cells ectopically expressing *ratA* displayed an even faster growth than the wildtype cells after an initial period of comparable slow growth. Though we did employ a low-copy plasmid, these results might still be indicative of a copy number effect. Nevertheless, these findings indicate that the lack of RatA results in an impaired, but not abolished ETC.

Since several studies have established that the plethora of phenotypes associated with the loss of Coq10 can be alleviated either by the addition of Q_2_ or by overexpression of the ubiquinone biosynthesis gene *coq8* as hallmarks of *coq10* deletion strains^17–20,27^, we investigated whether the small colony phenotype upon *ratA* deletion^13,21,23^ can equally be complemented by the addition of Q_2_ or by overexpressing *ubiB*, the bacterial homologue to *coq8*. As shown in Figure 5A, we observed a marginal, but statistically significant increase in colony size upon addition of Q_2_. A more pronounced increase in colony size was observed upon overexpression of *ubiB*, though a complete complementation was not obtained (Figure 5B). In summary, these results further solidify that RatA and Coq10 are homologues required for the proper functionality of the aerobic ETC, while precise molecular functions remain elusive.

**Figure 5.**
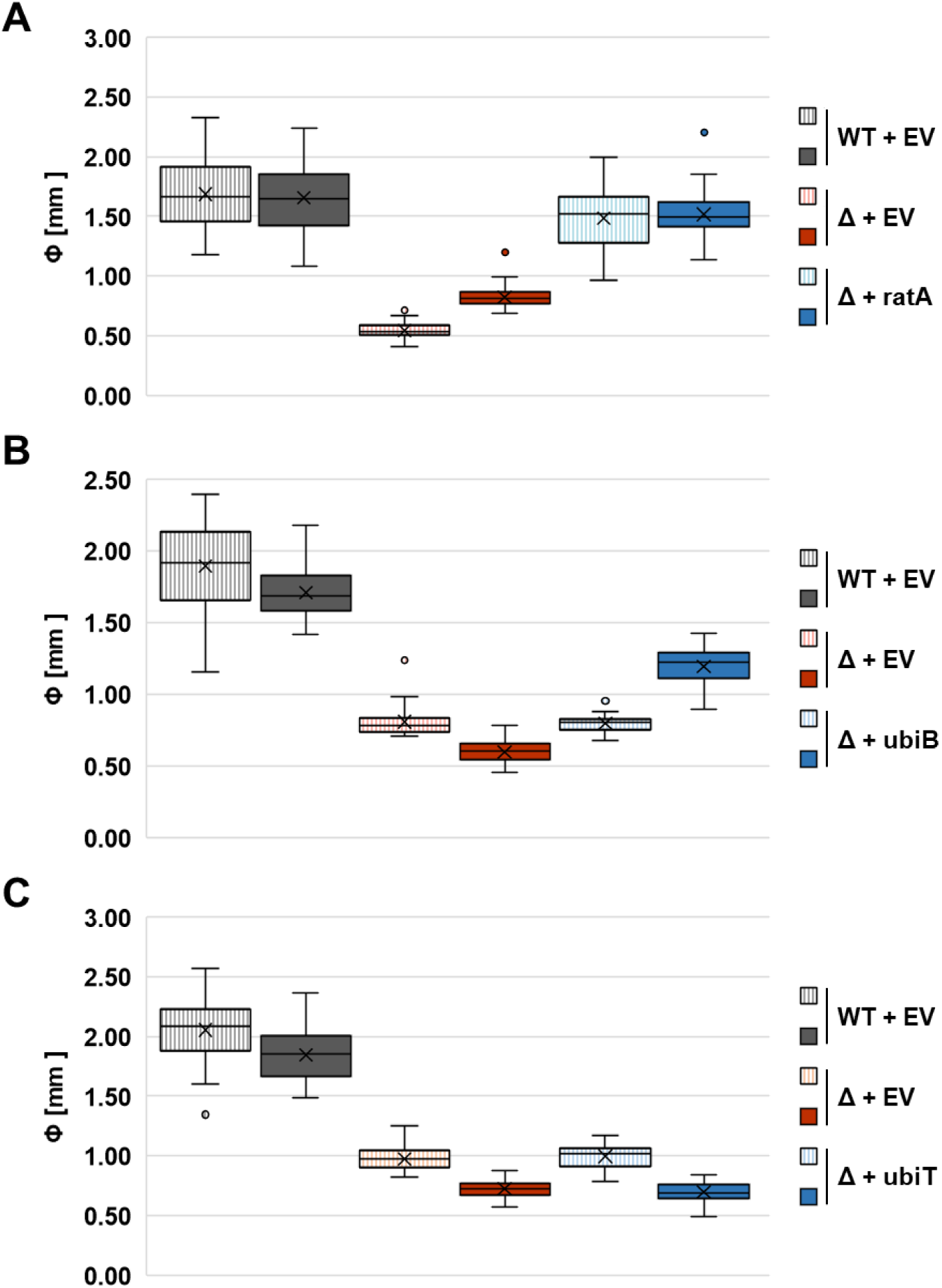
The small colony phenotype of a *ratA* deletion strain is partially rescued by the addition of Q_2_ and the overexpression of *ubiB*. Box and whisker plots representing colony sizes of a minimum of 30 individual colonies. Striped boxes represent experiments performed on LB plates with no additives, solid boxes represent experiments performed on LB plates supplemented with either 50 µM coenzyme Q_2_ (A) or 0.2% L-arabinose (B, C). **A** The *ratA* deletion strain transformed with the empty vector pZS*2 (Δ + EV, red) displayed a small colony phenotype compared to the corresponding wildtype *E. coli* strain BW25113 transformed with the same empty vector (WT + EV, grey). Ectopically expressing the *ratA* gene in the deletion strain under endogenous promoter control from the pZS*2-ratA-His plasmid (Δ + ratA, blue) mostly restored the observed phenotype. Furthermore, addition of Q_2_ partially rescued the small colony phenotype in the knockout strain. **B** Similar to A, the BW25113(Δ*ratA*) strain was transformed with pBAD-ubiB (Δ + ubiB, blue) for arabinose inducible overexpression of the gene. Addition of arabinose partially rescued the small colony phenotype of the *ratA* deletion strain transformed with the pBAD33 empty vector (Δ + EV, red) compared to the corresponding wildtype strain transformed with the same empty vector (WT + EV, grey). **C** Same as in B, but with the pBAD-ubiT plasmid transformed (Δ + ubiT, blue), which did not rescue the small colony phenotype after the addition of arabinose.

### The *ratA* deletion strain displays altered energy metabolism

Previous studies established that neither deletion of *coq10*, *pasT*, nor *ratA* resulted in drastically reduced total ubiquinone levels^15,17,21^. Fino et al. thus argued that the plethora of observed phenotypes upon loss of the lipid chaperon are likely indirect and downstream effects of broad distortions of bacterial physiology caused by a defective respiration apparatus^21^. We therefore investigated the cellular composition and metabolic state of a *ratA* deletion strain by performing proteomic and metabolomic analyses. As preliminary results revealed that deletion of the *ratA* gene simultaneously resulted in the strongly reduced expression of the downstream *ratB* gene, which overlaps with the end of the *ratA* ORF (Supplementary Figure 1A), we decided to employ a BW25113(Δ*ratAB*) strain. This strain was either complemented with a low-copy plasmid carrying the *ratAB* operon under endogenous promoter control or transformed with the corresponding empty vector.

Further validating this approach, previous studies showed that the loss of RatA, not RatB, was responsible for the difference in growth^22^. Both strains were grown until mid-exponential phase, at which point cells were simultaneously processed for both proteomics and metabolomics analysis. Finally, all identified metabolites and proteins were fed into the joint pathway analysis tool from Metaboanalyst^28^. As shown in Figure 6, we observed broad adaptations of several pathways of the central energy metabolism, including in the TCA cycle in response to the lack of the *ratAB* operon. As shown in Supplementary Figure 6, the mapped features, i.e. the identified gene and metabolites belonging to a specific pathway, could be both up- and/or downregulated (see also Supplementary Data 1 for more specifics on the mapped features). Complementary to the proposed role of RatA as a lipid chaperone for Q_8_, we observed only downregulations for the features in the oxidative phosphorylation pathway (Supplementary Figure 6). For the alanine, aspartate and glutamate metabolism, we observed both up- and downregulated features (Supplementary Figure 6). Since *E. coli* utilizes the amino acids provided in the LB medium in a sequential manner^26,29^, we hypothesized that the differential regulation of the metabolic pathways could alternatively be explained by the strains being at slightly different growth stages at the time of harvesting. Therefore, in order to further substantiate our findings, we queried a previously published dataset analyzing the metabolome of every available single gene mutant from the Keio collection for the influence of the *ratA* deletion on the metabolic state of the corresponding *E. coli* cells^30^. In contrast to our data, this work employed a chromatography-free system, which cannot resolve metabolites with similar molecular weight but, due to the genome wide approach, allows for the matching of the metabolome profiles of genes of unknown function to the profiles of well investigated genes. Fitting to our data, deletion of *ratA* (or *yfjG* in the dataset) resulted in adaptation of a variety of amino acids metabolisms including the alanine metabolism, of ABC transporters, and of the TCA cycle^30^. Interestingly, among the top five gene deletions with the highest similarities in their metabolome profile compared to the *ratA* deletion profile were the ubiquinone biosynthesis gene *ubiF*, the cytochrome bd-II subunit gene *appB*, and one of three superoxide dismutase genes *sodC*^30^. We therefore further focused our analysis on specific pathways and complexes of interest related to their function.

**Figure 6.**
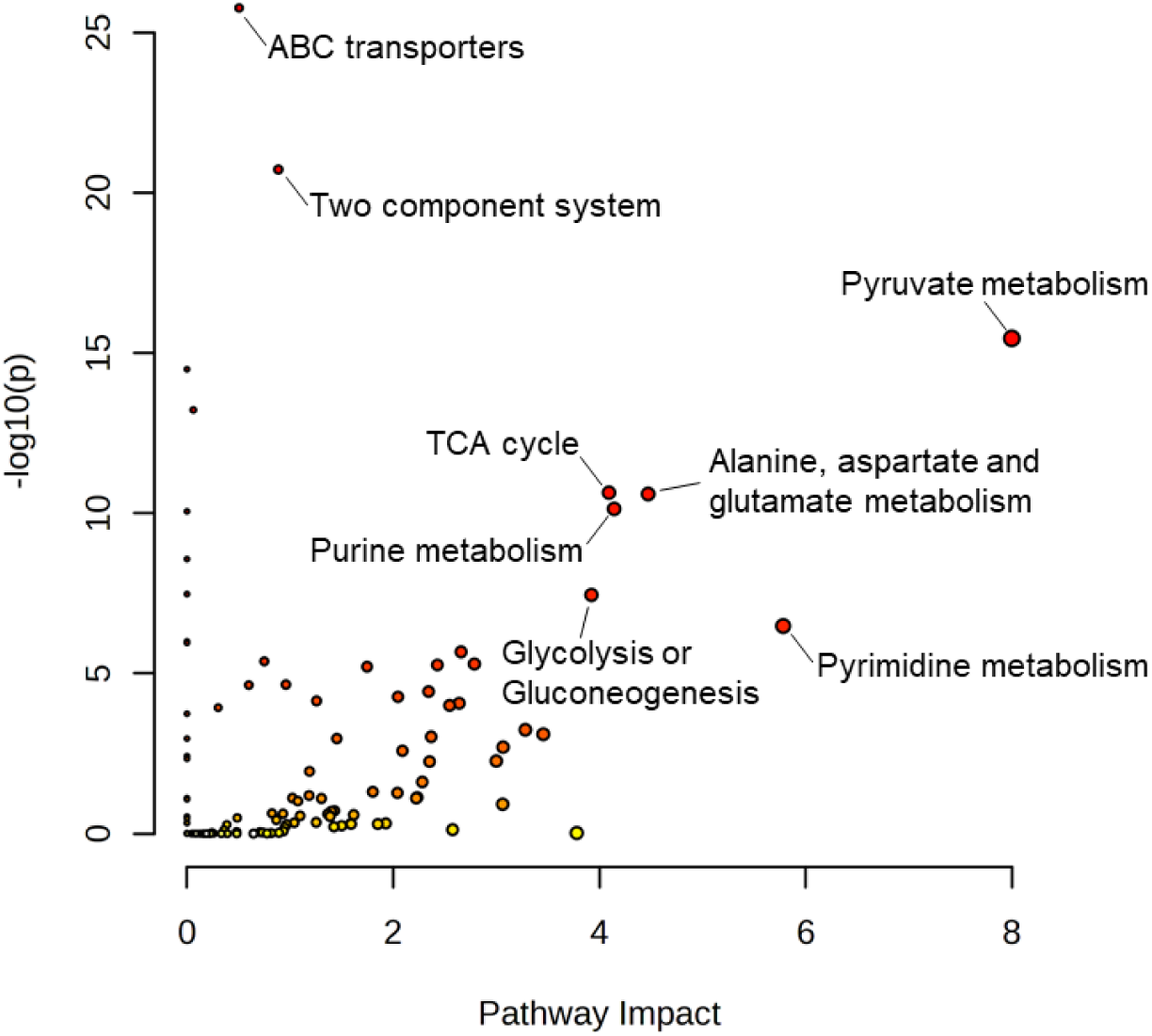
Loss of the *ratAB* operon broadly affects energy metabolism. Scatter plot showing the results of a joint pathway analysis performed on MetaboAnalyst^28^ with all the metabolites and proteins identified in the metabolomics and proteomics analysis. Displayed are all matched pathways according to the p values from the pathway enrichment analysis and pathway impact values from the pathway topology analysis.

### Aerobic ubiquinone biosynthesis pathway proteins are not affected by the loss of RatA

As both Coq10 and RatA have been proposed to be a lipid chaperone involved either in the synthesis of ubiquinone or in the correct positioning of ubiquinone in the ETC, we first focused on the proteins involved in the aerobic biosynthesis of Q_8_. Our proteomics data revealed no significant changes in the abundance of any of the involved enzymes (Supplementary Data 2). However, two recent studies established that *E. coli* cells are capable of performing Q_8_ synthesis even under anaerobic conditions^31,32^. To this end, a subset of the aerobic pathway enzymes is employed in combination with three additional enzymes termed UbiT, UbiU, and UbiV^31^. UbiU and UbiV perform O_2_-independent hydroxylation of the benzene ring reactions, whereas UbiT, which contains the same SCP2 domain as UbiJ, likely works as a scaffold for the anaerobic metabolon^32^. In the *ratA* mutant, we observed an upregulation of UbiT and downregulation of both UbiU and UbiV synthesis. Since both UbiT and RatA are lipid binding proteins, we speculated that the upregulation of UbiT could be a compensatory response to the absence of RatA. However, when we further overexpressed *ubiT* in the *ratA* deletion strain, we observed no increase in colony size (Figure 5C). We therefore suggest that the observed increase in UbiT concentration is not an actual regulatory response, but rather a product of the slightly different growth stages of the wildtype and the *ratA* deletion strain at the time of harvesting.

### Aerobic respiratory complexes remain largely unaffected by the absence of RatA

As previous studies in yeast established a negative impact on respiration upon loss of Coq10^15,17–19^, we further explored potential changes in the abundance of the aerobic respiratory complexes in our proteomics dataset (Supplementary Data 2). As shown in Figure 1A, there are three complexes in the ETC of *E. coli* capable of reducing ubiquinone to ubiquinol - NADH:quinone oxidoreductase I (NDH-1), NADH:quinone oxidoreductase II (NDH-2), and succinate:quinone oxidoreductase (SDH). The concentrations of proteins building up the SDH and NDH-1 complexes were largely unaffected by the loss of RatA (log_2_ FC range of - 0.13 to 0.62), whereas the abundance of the singular NDH-2 protein was upregulated almost twofold compared to the wildtype (log_2_ FC of 0.96). After reduction, ubiquinol transports the electrons to one of the three major ubiquinol:oxygen oxidoreductases, which vary in abundance based on oxygen availability^33^. We observed no drastic change in abundance of the detected subunits of cytochrome bo_3_ in our *ratA* deletion strain (log_2_ FC range of 0.31 to 0.46), whereas both the cytochromes bd-I and bd-II were potentially slightly downregulated (log_2_ FC range of -0.61 to -1.13).

NDH-1 and cytochrome bo_3_ are both proton pumping complexes, while cytochromes bd-I and bd-II do not pump additional protons across the inner membrane and contribute to the formation of a proton motive force (PMF) solely by a vectorial charge transfer^34^. In contrast, NDH-2 does not contribute to the generation of the proton potential in either way^35^. Consequently, an upregulation of NDH-2 likely contributes to the lower membrane potential previously reported upon deletion of the *pasTI* and *ratAB* operon^21^. To confirm these findings, we investigated the motility of *E. coli* cells in the presence or absence of RatA. As the rotary motion of *E. coli* flagella is driven by proton flux from the periplasm into the cytosol^36,37^, motility can serve as a proxy to measure the PMF. As shown in Figure 7, we observed a significant reduction in the swimming radius upon loss of RatA. In contrast, despite the majority of ATP being synthesized by the PMF-fueled ATP synthase, our metabolomics data revealed no change in ATP abundance between the wildtype and *ratA* deletion strain (log_2_ FC of 0.09, Supplementary Data 1).

**Figure 7.**
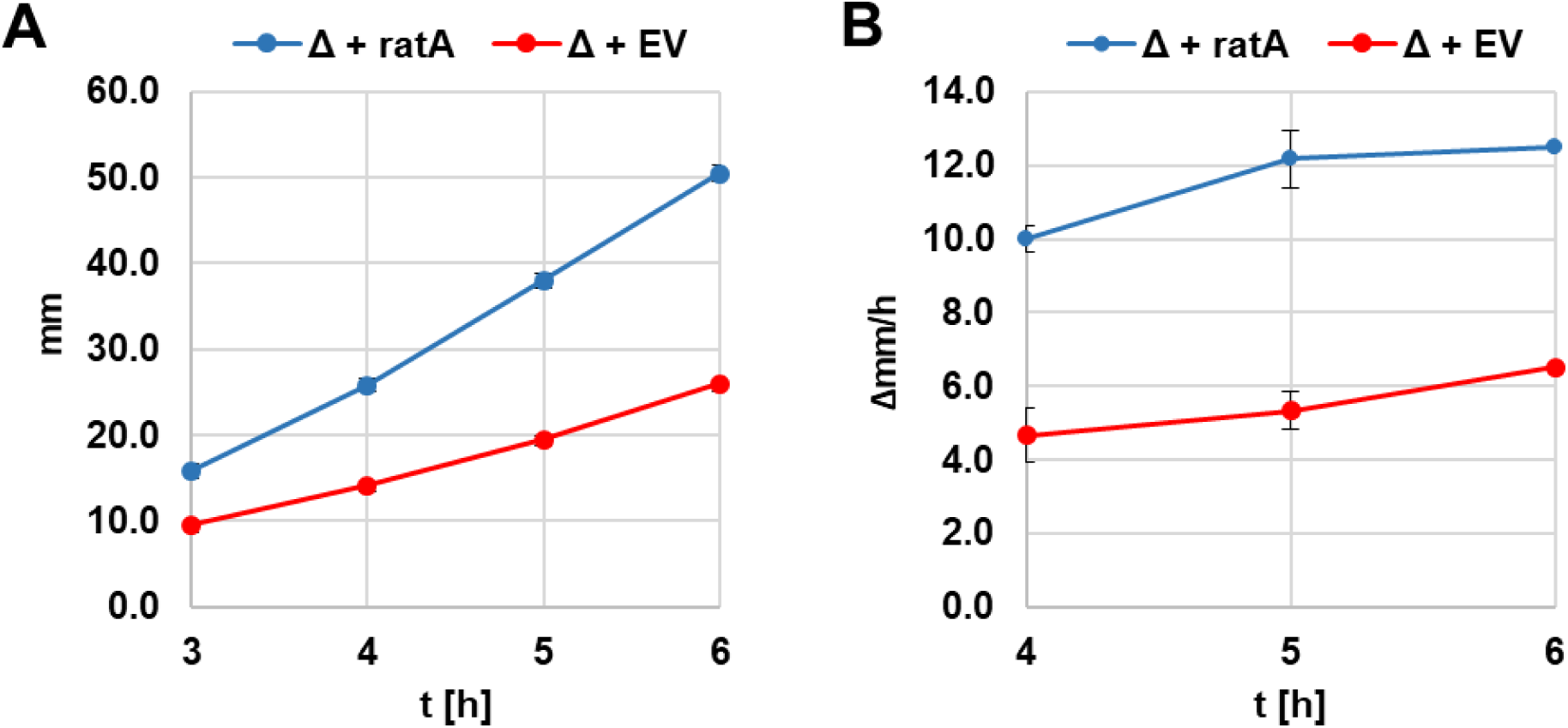
Loss of RatA negatively influences bacterial motility. The motility of BW25113(Δ*ratA*) transformed either with the pZS*2-ratA-His plasmid ectopically expressing the *ratA* gene under endogenous promoter control (Δ + ratA, blue line) or the corresponding empty vector pZS*2 (Δ + EV, red line) was assessed by measuring the swimming radius of the cells incubated in LB swimming plates at 37°C. **A** Average absolute values measured at the indicated time points with error bars representing the standard deviation (n=3). **B** Average change in radius per hour calculated by subtracting the previously measured radius from the current radius at the indicated time point (n=3).

### The *ratA* deletion strain displays signs of increased oxidative stress

Previous studies on *coq10* and *pasT/ratA* deletion strains demonstrated an increase of oxidative stress symptoms^13,16,18,20,21,38^. While it has been shown that respiratory complexes are not the major source of ROS in *E. coli*, both *in vitro* and *in vivo* studies established that NDH-2 contributes to their formation under aerobic growth conditions^39,40^. We thus screened our proteomics and metabolomics data for signs of increased oxidative stress upon *ratA* deletion. In *E. coli*, depending on the ROS species, either the OxyR or SoxRS transcription factors are activated to mitigate the damage of oxidative stress^41^. OxyR is directly activated by increased H_2_O_2_ levels. We observed no difference in OxyR abundance (log_2_ FC of -0.11) and no clear upregulation of the proteins in the OxyR regulon (Supplementary Data 2). In fact, the major H_2_O_2_ scavenging catalase-peroxidase KatG was downregulated in the *ratA* deletion strain (log_2_ FC of -1.64), indicating that H_2_O_2_ increase was not a source of oxidative stress. The SoxRS system relies on SoxR, which is constantly present, first being activated by superoxide (O_2_^-^) through the oxidation of bound iron-sulfur clusters on the protein, thus allowing it to induce the expression of *soxS*^41^. Fittingly, the abundance of SoxR was not affected (log_2_ FC of 0.13), but a strong upregulation of the transcription factor SoxS was detected (log_2_ FC of 2.71). Furthermore, we observed the upregulation of several proteins of the SoxS regulon, including the superoxide dismutase SodA and the fumarase isozyme FumC. FumC is one of three characterized fumarase isozymes in *E. coli* that catalyze the hydration of fumarate to L-malate in the TCA cycle. Previous work has shown that unlike FumA and FumB, FumC does not require iron as a co-factor for its activity, making it less susceptible to O_2_^-^ and allowing it to serve as a backup enzyme during superoxide stress conditions^42^. Thus, in summary, several lines of evidence indicate increased oxidative stress based on elevated levels of O_2_^-^ in absence of RatA.

### RatA interacts with proteins in the ubiquinone biosynthesis metabolon

Coq10 is an inner mitochondrial membrane protein^15,17^, while the physiologically relevant form of RatA, RatA_Δ14_, contains no predicted membrane domain. We thus investigated the localization of RatA by separating *E. coli* cells into cytosolic and membrane fractions and analyzed the presence of RatA in the respective compartments by western blot analysis. As shown in Figure 8, RatA is detected in both the cytosolic and membrane fraction.

**Figure 8.**
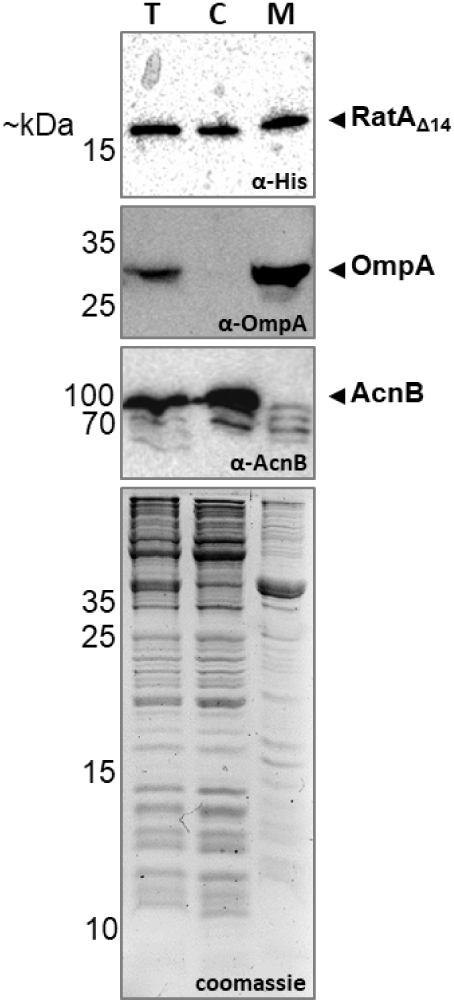
RatA is present in both the cytosol and the membrane. Western blot analysis of samples harvested from the BW25113(Δ*ratA*) pZS*2–ratA–His strain ectopically expressing the wildtype *ratA* gene with a 3′-terminal His-tag sequence under endogenous promoter control to investigate the cellular localization of the RatA protein. The total cell extract (T) was fractionated into a cytosolic (C) and membrane fraction (M). As fractionation controls, the outer membrane protein OmpA and the cytosolic aconitase AcnB were used in combination with a Coomassie gel staining the respective proteomes of each fraction.

As described above, in *E. coli*, Q_8_ synthesis begins in the inner membrane and is finalized in a large cytosolic metabolon^10^. However, it remains unknown how the hydrophobic ubiquinone precursor molecules are delivered from the membrane to the metabolon and how the mature Q_8_ is transported back to the inner membrane. Therefore, based on its localization, we hypothesized that RatA performs a shuttling function between the metabolon and the inner membrane. To this end, we performed bacterial two hybrid (BACTH) assays to investigate potential interactions of RatA with any of the Ubi proteins. As shown in Figure 9, we observed a clear interaction of RatA with enzymes UbiF and UbiG. Interestingly, these two enzymes are performing the two final steps of Q_8_ biosynthesis: UbiF hydroxylates the C-5 of the benzene ring^43^, which is then further O-methylated by UbiG to form the mature ubiquinone^44^. This is in line with our previous finding that RatA binds mature Q_8_ (Figure 3). Therefore, combining these results, we propose that RatA is responsible for the shuttling of newly synthesized Q_8_ from the metabolon to the membrane.

**Figure 9.**
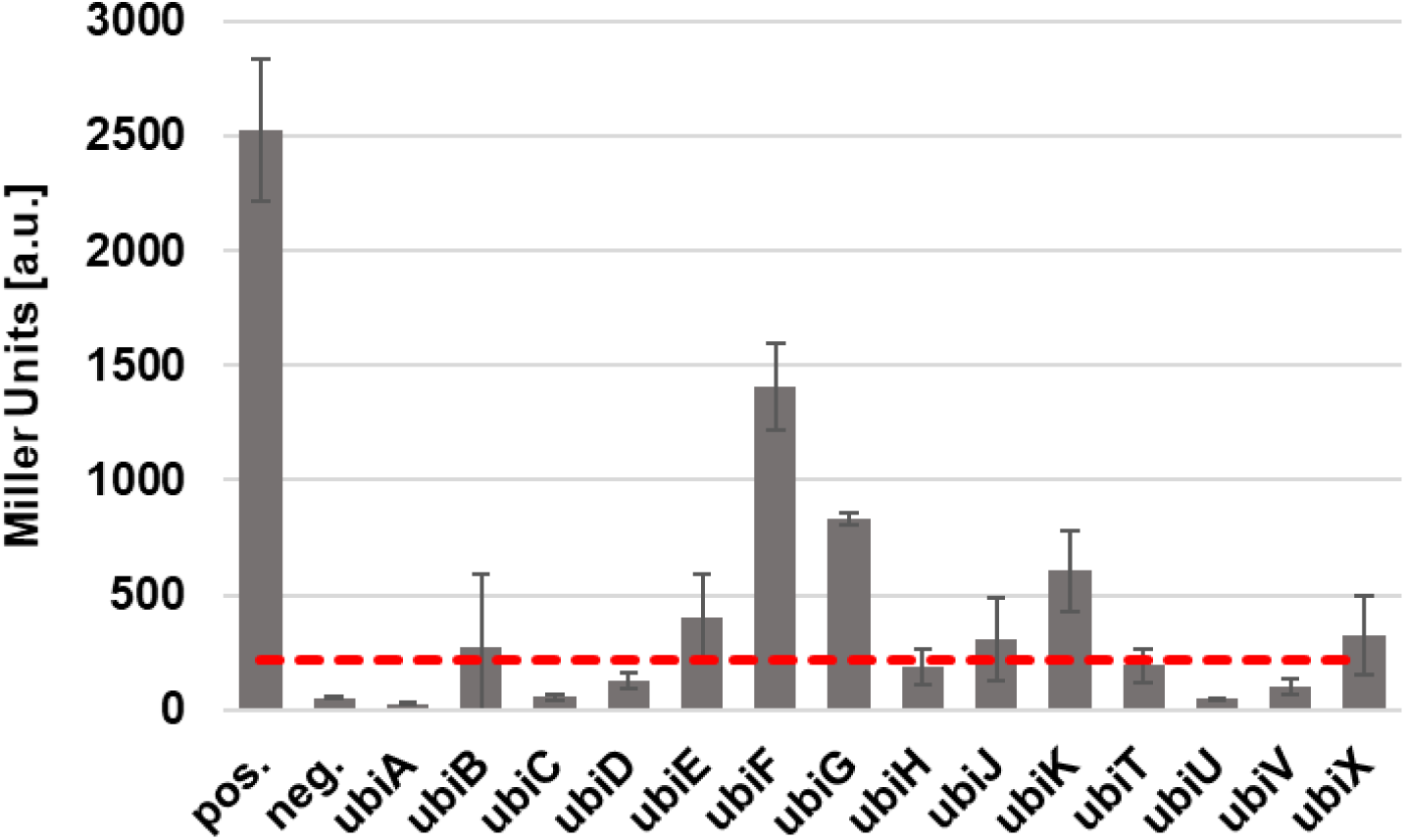
RatA interacts with ubiquinone biosynthesis proteins. Protein-protein interactions were assessed by employing Bacterial Two-Hybrid (BACTH) assays in the reporter strain BTH101. For each assay, the plasmid pKNT25-ratA_Δ14_ was transformed in combination with the plasmid pUT18C carrying the indicated *ubi* genes. As a positive control, the pKT25-zip plasmid was transformed in combination with the pUT18-zip plasmid. As a negative control, plasmids pKNT25 and pUT18C were employed. For quantification, β-galactosidase assays were performed in triplicates, with the red dotted line representing four times the average value of the negative control.

## Discussion

### RatA is not a toxin

RatA was initially described to target the 50S ribosomal subunit preventing translation initiation, thus being the toxin-moiety of the proposed RatAB TA pair^11,12^. A later study performed in uropathogenic *E. coli* revealed both a toxic and a beneficial stress resistance role of RatA depending on its expression level^13^, thus the module was termed PasTI, for persistence and stress-resistance toxin and immunity proteins. Moreover, the toxic effect was only observed upon overexpression of the full-length *pasT* gene and was alleviated by simultaneous overexpression of *pasI*^13^. In contrast, the stress resistance effects were observed upon low-level expression of *pasT* and tolerated the deletion of the sequence encoding for the first 13 amino acids of the protein^13^. In a third study by Fino et al. aimed at investigating in more detail the biological function of RatAB/PasTI in *E. coli*, the authors disputed the notion that PasT is part of a bona-fide TA pair and instead provided evidence that PasT is the bacterial homologue of the mitochondrial protein Coq10^14^. We recently identified that these contradictory results can be attributed to a mis-annotation of the gene in the reference genome^23^. We showed that overexpression of the coding sequence of *ratA* as currently annotated in the NCBI reference sequence NC_000913.3 results in the synthesis of two RatA variants with each one exhibiting the different effects on the *E. coli* cells as previously observed. Overproduction of the full-length RatA (RatA_fl_) originating from the annotated start codon (Met1) was toxic, whereas the shorter RatA_Δ14_ originating from the Met14 codon was beneficial and could alleviate the small colony phenotype observed upon deletion of the *ratA* gene^23^.

Furthermore, we identified an endogenous σ^70^ promoter for *ratA* that is partially located inside of the annotated ORF, resulting in a transcription start side downstream of the annotated start codon and thereby excluding the possibility of toxic RatA_fl_ formation^23^. However, the presence of alternative promoters further upstream that could give rise to production of the toxic RatA_fl_ protein cannot be excluded. Therefore, in this study, we investigated whether RatA_fl_ is synthesized under conditions previously reported to result in a detectable phenotype upon deletion of the *ratA* gene from the genome. As shown in Figure 2, only the shorter, non-toxic RatA_Δ14_ was detectable independent of the stress condition. In combination with the observation that the deletion mutant of the putative anti-toxin *ratB* is viable and taking the strong similarities between RatA and the mitochondrial protein Coq10 into account, we ultimately conclude that the only physiologically relevant RatA variant in *E. coli* is RatA_Δ14_ and its role is not that of a toxin.

### The function of RatB remains elusive

RatB was suggested to be the putative anti-toxin based on its genomic location with the start of its ORF overlapping with the end of the *ratA* gene (Supplementary Figure 1A). Consequently, we observed that deletion of *ratA* alone results in the simultaneous strong downregulation of *ratB* expression (log_2_ FC of -6.75 in data not shown). Despite this strong genetic connection, in the original work describing RatA as a ribosome-targeting toxin, the authors reported that they failed to achieve complex formation between purified RatA and RatB and did not observe a neutralizing effect on the toxicity of RatA when *ratB* was co-expressed on plates^12^. In contrast, co-overexpression of *pasI* with *pasT* in liquid culture did alleviate the toxic effect^13^. Therefore, while RatB is definitionally not an anti-toxin to the non-toxic RatA, it nevertheless appears to be able to regulate RatA activity under specific conditions. However, the exact function of RatB remains elusive. A protein BLAST^45^ approach reveals that RatB is a RnfH family protein, which are usually either encoded in an operon with a group of *rnf* genes responsible for the formation of a Na^+^ coupled respiratory complex^46,47^ or as part of the *ratAB* operon^48^. As Fino et al. have noted, *ratAB/pasTI* operons are only commonly found in beta-and gammaproteobacteria, whereas solitary *ratA/pasT* genes are generally much more abundant^14^. Consistent with this observation, the variety of phenotypes observed upon deletion of the *ratAB* operon could always be complemented by the re-introduction of RatA alone^13,14,22,23^, hinting more to a fine-tuning role of RatB. To investigate the function of RatB in greater detail, during this work, we employed several different approaches to visualize the protein by western blot analysis. We aimed to produce N- or C-terminally tagged RatB protein using different plasmids systems and either its endogenous promoter or a variety of standard, metabolite-inducible promoters. Unfortunately, we were not able to detect the protein with any tested approach (data not shown), suggesting that the RatB protein does not easily tolerate manipulation.

### RatA, the bacterial Coq10 homologue, influences the central energy metabolism

Previously, Fino et al. determined RatA to be the bacterial homologue of the mitochondrial protein Coq10 based on the following observations: First, both proteins contain a START domain. Second, as previously observed for *coq10* mutants, deletion of *ratA* results in a plethora of phenotypes indicative of a defective ubiquinone-dependent ETC. Third, such phenotypes observed in bacteria could largely be complemented by human Coq10. In this study, we further substantiated the notion that RatA is the bacterial equivalent of Coq10 by exploring two additional hallmarks of *coq10* mutants in our *ratA* deletion strain. As shown in Figure 5A and B, both the addition of Q_2_ and the overexpression of *ubiB*, the bacterial equivalent of *coq8*, could partially rescue the small colony phenotype of a *ratA* deletion strain, corroborating previous studies on Coq10^17–20,27^. It is still unclear how UbiB contributes to Q_8_ synthesis. While it was suggested that UbiB is responsible for the C5-hydroxylation, later work showed that this reaction was in fact performed by UbiI^49^. On the other hand, Coq8 has been proposed to be a protein kinase targeting Coq3^50^. UbiB might therefore equally be a kinase as initially proposed^51^, targeting the bacterial equivalent of Coq3, the later discussed UbiG.

If the observed phenotypes in the *ratA* deletion strain are indeed the result of a defect in the ubiquinone-dependent ETC, we expect that growth of RatA-deficient cells is impaired on the TCA cycle intermediate succinate as the sole carbon source compared to glucose, the preferred and fermentable carbon source of *E. coli*. As shown in Figure 4C and D, this notion held true. Importantly, while growth was demonstrably affected, compared to the *ubiGJ* double deletion strain that is completely lacking Q_8_, loss of RatA did not result in cells that were completely incapable of growing on succinate. Combining these findings with both the previously assessed reduction of the membrane potential^14^ and with the observed reduction in PMF-driven motility in the *ratA* deletion strain (Figure 7), we concluded that the absence of RatA results in an impaired, but not abolished ETC. The adapted energy metabolism of the *ratA* deletion cells that is likely contributing to a superoxide-driven increase in oxidative stress is discussed in detail above. It should however be noted, that we did not observe a change in ATP abundance between the parental and deletion strain. We speculate that either the PMF is preferentially utilized for ATP production before motility or that this observation is indicative for the robustness of energy production in *E. coli* by a variety of different pathways.

Contrary to previously published results^13^, we observed that the impaired ETC results in a reduced growth phenotype even in nutrient rich LB medium (Figure 4A, B). Interestingly, this growth phenotype was culture volume dependent (Supplementary Figure 4), suggesting that the decreased volume to surface ratio decreases aeration, forcing the cells to adapt to micro-aerobic conditions, which diminishes the growth phenotype similar to a previous observation for the small colony phenotype of a *ratA* deletion strain under anaerobic growth conditions^21^.

### RatA is a shuttle for ubiquinone between its site of synthesis and the inner membrane

While the affected respiration of *ratA* deficient strains can potentially explain the plethora of phenotypes observed (e.g. respiratory efficacy was previously shown to influence persister cell formation^52^), the precise molecular function of RatA remained elusive. Like its mitochondrial homologue Coq10, RatA was proposed to serve a lipid chaperone function either facilitating *de novo* ubiquinone biosynthesis, transporting ubiquinone from the metabolon to its final position in the ETC, and/or assisting the shuttling of ubiquinone between the different complexes of the ETC. However, these proposed functions relied on the assumption that RatA indeed binds Q_8_. While both the structure and sequence conservation between RatA and Coq10 are certainly a valid indication, binding of Q_8_ to RatA was previously not confirmed. In this work, we filled this experimental gap and demonstrated Q_8_ binding to RatA (Figure 3). Our studies to identify residues important for Q_8_ binding revealed that mutations in the putative binding cavity strongly affected protein abundance (Supplementary Figure 3C, D). Consequently, phenotypes observed upon mutation of such residues should generally be carefully analyzed by simultaneously accounting for the protein level.

To further investigate the precise function of RatA, we first determined its cellular location. In contrast to Coq10, which was shown to be a uniformly distributed inner mitochondrial membrane protein^8,15,17^, RatA is present in both the cytosolic and in the membrane fraction (Figure 8). Previously it was shown that in *E. coli*, Q_8_ synthesis begins in the inner membrane and is finalized by an approximately 1 MDa complex in the cytosol^10^. However, it remains unclear how the ubiquinone precursors are transported from the initial complexes in the membrane to the cytosolic complex and how the mature Q_8_ is transported from this complex back to the inner membrane to its functional position in the ETC. Employing a BACTH approach, we thus investigated whether RatA interacts with the proteins of the Q_8_ synthesis machinery. Most consistently, we observed interactions between RatA and UbiG and UbiF (Figure 9), which catalyze the last two steps in the ubiquinone biosynthesis. While we did potentially observe a slight interaction of RatA with the early synthesis proteins UbiB and UbiX, these results were less consistent and less pronounced. We thus propose that RatA serves as a shuttle for Q_8_ between the cytosolic metabolon and the inner membrane. What remains to be elucidated is whether RatA transports the Q_8_ to a specific protein (complex) of the ETC or whether it deposits its lipid load directly into the inner membrane. Previously, Fino et al. showed that the observed phenotypes upon loss of RatA can be largely complement by human Coq10 and thus anticipated that *E. coli* systems could be used for future studies to unravel the apparently conserved molecular function of RatA/Coq10 proteins^14^. Based on the RatA localization results presented here, we would again caution to directly translate our findings to eukaryotic system.

In summary, we have shown that RatA is not a toxin, confirmed and extended on previously published results highlighting similarities between Coq10 and RatA, proved for the first time that RatA can bind Q_8_, and established that RatA can interact with the ubiquinone biosynthesis metabolon. We thus propose to unify the naming of RatA and PasT to UbiM (ubiquinone Metro) to *i)* reflect its function and *ii)* to be in accordance with the naming convention of the other ubiquinone biosynthesis proteins.

## Material and Methods

### Bacterial strains and plasmid construction

Wildtype *E. coli* K-12 strain BW25113 was obtained from the Keio collection^53^ and the corresponding *ratA* deletion strain complemented with a low-copy plasmid carrying *ratA* under endogenous promoter control has been described before^23^ (see also Supplementary Figure 1). The closely related uropathogenic *E. coli* strain CFT073 was a kind gift from the laboratory of Prof. Karin Schnetz (Universität zu Köln). Removal of the *ratAB* or *pasTI* operon from the BW25113 or CFT073 genome and excision of the kanamycin resistance cassette using the heat sensitive plasmid pCP20 were performed according to the original publication by Datsenko and Wanner^54^.

For detailed information on the plasmids employed in this study and for the cloning of the individual plasmids, see Supplementary Data 3. Standard protocol of cloning involved the amplification of genomic regions of interest by PCR (Thermo Scientific, 2x Phusion Master Mix with HF Buffer) followed by digestion with the appropriate FastDigest restriction enzymes from Thermo Scientific. Digested fragments were ligated into the corresponding, equally digested vectors employing the T4 DNA Ligase from Thermo Scientific before transformation into competent Top10 cells for plasmid amplification. Point mutations were introduced by inverse PCR followed by blunt-end ligation. Finally, plasmids were isolated with the innuPREP Plasmid Mini Kit 2.0 (iST Innuscreen GMBH) and sequenced by automated Sanger sequencing (Mycrosynth).

#### General bacterial culturing

For long term storage, constructed strains were stored in 25% glycerol solutions at -80°C. If required for plasmid retention, all described media and plates were supplemented with either 25 µg/ml kanamycin, 30 µg/ml chloramphenicol, and/or 100 µg/ml ampicillin. For biological replicates, the glycerol stocks were routinely streaked out on LB plates (10 g/L peptone, 5 g/L yeast extract, 10 g/L NaCl, 1.5% w/v agar) and incubated overnight at 37°C before selecting single colonies from the plate to inoculate either LB medium or 1x M9+glucose pre-cultures (1x M9 salt mix (47.4 mM Na_2_HPO_4_, 22 mM KH_2_PO_4_, 8.5 mM NaCl, 18.7 mM NH_4_Cl, adjusted to pH 7),1 mM MgSO_4_, 0.1 mM CaCl_2_, 0.2% w/v glucose). The pre-cultures were routinely grown overnight at 37°C, 165 rpm before re-dilution in fresh medium to a starting OD_600_ of 0.05. If grown in flasks, cultures were standardly grown at 37°C, 165 rpm and growth was monitored by measuring the OD_600_. For plate reader experiments, if not specified otherwise,100 µl of the cultures were added per well in a 96 well plate (Thermo Scientific, #612F96, flat bottom). Generally, two technical replicates were measured per biological replicate. Plates were incubated in the BioTek Synergy H1 Microplate Reader at 37°C while continuously performing a fast orbital shake with an orbital frequency of 355 cpm (4 mm) and measuring the A_600_ every 15 minutes.

#### Influence of the carbon source

For growth with different carbon sources, cells were pre-cultured overnight in 1x M9+glucose medium and re-diluted the next day in 1x M9 with no carbon source added. The re-diluted culture was further split into smaller aliquots and either glucose or succinate were added to a final concentration of 0.2% (w/v) or 0.3% (w/v), respectively. For the nutritional stress assay, the strains were pre-incubated overnight and re-diluted the next day in an adapted 1x M9 medium containing 0.1% (w/v) glucose, 0.2% (w/v) casamino acids (CAS), and 16.5 µg/ml thiamin to mimic previously applied conditions^13^. Cultures were incubated at 37°C, 165 rpm to an OD_600_ of approximately 0.4, when cells were pelleted by centrifugation at 3,000 g for 5 minutes at room temperature before being resuspended in an equal volume of prewarmed 1x M9 medium with no additional CAS added. Afterwards, the cultures were continued to be incubated at 37°C, 165 rpm while monitoring the OD_600_. Samples for western blot analysis were harvested as described below.

#### Influence of different stress conditions

For oxidative stress and antibiotic treatment conditions, cells were both pre-cultured overnight and re-diluted the next day in LB medium. The cultures were incubated at 37°C, 165 rpm until they reached an OD_600_ of approximately 0.4, when either paraquat (i.e. methyl viologen, MV) was added to a f.c. of 1 mM, hydrogen peroxide (H_2_O_2_) was added to a f.c. of 10 mM, ampicillin was added to a f.c. of 100 µg/ml, or ciprofloxacin was added to a f.c. of 10 µg/ml. H_2_O_2_ treated cells were further incubated for 30 minutes, MV treated cells for 1 hour, and antibiotic treated cells for 5 hours. Samples for western blot analysis were harvested as described below.

#### Western blot

At indicated time points, the OD_600_ of the bacterial culture was measured and samples were harvested at 10,000 g for 2 minutes at room temperature. The resulting pellet was resuspended in 2x Laemmli buffer (4% SDS, 20% glycerol, 0.004% bromophenol blue, 125 mM Tris base, 10% β-mercaptoethanol, pH 6.8) to a final calculated concentration of 20 OD units per mL and incubated at 85°C for 5 minutes while shaking at 1,000 rpm. On 16% tris-tricine gels, 0.1 OD units were loaded per sample and separated proteins were wet transferred to 0.2 μm nitrocellulose membranes (Amersham Protran, Cytiva) at 100V for 1 hour at 4°C in 1x transfer buffer (25 mM Tris base, 192 mM Glycine, pH 8.3). Membranes were blocked with 5% (w/v) milk powder in 1x PBS supplemented with 1% Tween-20 (1x PBS-T) before addition of the following primary antibodies for incubation overnight at 4°C while shaking: anti-His (Cytiva, 27471001, raised in rabbit, 1:10,000 dilution), anti-OmpA (Antibody Research Corporation, #111120, raised in rabbit, 1:5,000 dilution), and anti-AcnB (gift from Jeff Green, University of Sheffield, raised in rabbit, 1:10,000 dilution). The horseradish peroxidase coupled anti-rabbit IgG (Cell Signaling Technology, 7074S) was used as a secondary antibody at a dilution of 1:10,000. All blots were developed using the SuperSignal West Pico PLUS Kit chemiluminescent reagent from Thermo Scientific in the BioRad ChemiDoc Imaging system.

#### Ubiquinone binding to RatA

For protein purification via a strep-tag, the wildtype BW25113 strain was transformed either with the pBAD-ratA_Δ13_-strep plasmid or the pBAD-hns-strep plasmid. Both strains were grown in LB medium at 37°C while shaking at 165 rpm to an OD_600_ of approximately 0.5 before induction with L-arabinose to a final concentration of 0.2 % (w/v). After induction, the cultures were incubated at 37°C, 165 rpm for two additional hours (OD_600_ ∼ 2) before harvesting the cells of 400 ml cultures at 4,000 rpm for 20 minutes at 4°C. All further proceedings were performed on ice and with cold buffers. Pellets were washed with 25 ml of Buffer W (100 mM Tris-HCl pH 8.0, 150 mM NaCl, 1 mM EDTA) and pelleted again at 4,000 rpm for 15 minutes at 4°C before finally being resuspended in 5 ml of Buffer W. Cells were disrupted at 1.9 kbar in the OneShot cell disruptor from Constant Systems Ltd. The resulting lysate was supplemented with 5 µl DNase I (10 U/µl) and incubated at 37°C for 5 minutes before addition of Triton-X100 to a final concentration of 0.1% (v/v). The lysates were first cleared by two sequential centrifugation steps at 4,000 rpm for 20 minutes at 4°C, followed by centrifugation at 30,000 g for 1 hour at 4°C. The cleared lysate was loaded onto 400 µl column bed volume (CBV) of high capacity Strep-Tactin XT Superflow beads from iba Life Sciences that were equilibrated beforehand by washing the beads two times with 1 CBV of Buffer W. Beads and lysates were incubated with rotation at 4°C for 1.5 hours. Unbound proteins were collected in the flow-through and the beads were washed first two times with 1 CBV of Buffer TX+ (Buffer W with 0.15 Triton-X100 and 40 mM DTT) and then five times with 1 CBV of Buffer W. Proteins were eluted with Buffer E (Buffer W with 2.5 mM D-desthiobiotin) containing 0.25 µg of coenzyme Q_2_ per 10 ml of elution buffer as a spike-in control. Eluted proteins were analyzed by western blot and 175 µl of eluted fractions were further processed with chloroform/methanol extraction for analysis of Q_8_ binding. 100 µl of the lower phase was diluted with 50 µl methanol and 1 µl of each extract was directly injected onto a Kinetex (Phenomenex) C8 column (100 Å, 100 x 2.1 mm), employing a 9-minute-long linear gradient from 90% A (60% acetonitrile, 39.8% water, 0.2% formic acid and 20 mM ammonium acetate) to 95% B (90% isopropanol, 5% acetonitrile, 4.8% water, 0.2% formic acid and 20 mM ammonium acetate) at a flow rate of 100 µl/min. Samples were analysed using precursor ion scanning by LC-MS/MS (liquid chromatography tandem mass spectrometry), employing the precursor ion scanning mode of a TSQ Altis mass spectrometer (Thermo Fisher Scientific) using the fragment ion 197.1 *m/z* characteristic for the coenzymes. Subsequently. they were detected, confirmed and quantified by LC-MS/MS, employing the selected reaction monitoring (SRM) mode of the mass spectrometer, using the transitions 727.5 *m/z* to 197.1 *m/z* (coenzyme Q_8_), and 319.2 *m/z* to 197.1 *m/z* (coenzyme Q_2_) in the positive ion mode. Authentic standards of the coenzymes have been used for determining optimal collision energies of the SRM transitions and for validating experimental retention times.

#### Colony size assay

For colony size determination, the different strains were pre-cultured and re-diluted in fresh LB medium to an OD_600_ of 0.05 as described above before further serial dilution in LB to a final dilution factor of 10^−5^. From the final dilution, 50 µl were plated on LB plates containing the required antibiotics for plasmid retention and either 50 µM of coenzyme Q2 or 0.2% (w/v) arabinose. For the negative controls, equal volumes of solvent (acetone for coenzyme Q2, water for arabinose) were added to the LB+agar solutions before plating. After plating, the colonies were developed at 37°C for 20 hours and individual colony sizes were measured in Fiji^55^.

#### Non-targeted proteomics and metabolomics for joint pathway analysis

100 ml bacterial cultures in LB medium were prepared as described above for the BW25113(Δ*ratAB*) strain transformed either with the low-copy plasmid pZS*2-ratAB encoding for the *ratAB* operon under endogenous promoter control or with the corresponding empty vector (pZS*2). The bacteria were grown for four hours at 37°C while shaking at 165 rpm, before dividing the cultures into two times 50 ml and harvesting the cells at 5,000 g for 5 minutes at room temperature. One pellet was further processed for proteomics analysis, whereas the other was further processed for non-targeted metabolomics analysis. In both cases, further proceedings were performed on ice.

#### Proteomics

For proteomics, the pellet was washed with 1 ml of ice-cold 1x PBS and pelleted again at 10,000 g for 1 minute at 4°C before the resulting pellet was finally resuspended in 1 ml of ice-cold 1x PBS. From this suspension, 30-40 µl were transferred to a fresh tube to account for slight differences in the original OD_600_ before harvesting. Cells were again pelleted at 10,000 g for 1 minute at 4°C before being resuspended in 100 µl hot lysis buffer (4% SDS, 100mM Tris HCl pH 8.5) and incubated at 95°C for 5 minutes while shaking at 1,000 rpm. To complete cell lysis, samples were sonicated in the Bioruptor (Diagenode) with 5 cycles of 30/30 seconds with high energy. Persisting viscosity was reduced using a sonication rod by applying 16 x 0.5 seconds pulses at 50% amplitude. Samples were centrifuged for 10 minutes at 20,000 g to pellet debris. Supernatants were reduced by adding DTT at a final concentration of 10 mM incubating for 30 minutes at room temperature. Free cysteines were alkylated with iodoacetamide at a final concentration of 20 mM, incubating for 30 minutes at room temperature in the dark. One hundred µg protein in 40 µL were mixed with Sera-MagSpeed Beads 1:1 mix (GE Life Sciences, cat. nos. 45152105050350 and 65152105050350) at a beads:protein ratio of 10:1. Ethanol was added to a final concentration of 50% and mixed thoroughly. The tubes were kept in a thermomixer at room temperature for 10 minutes at 1000 rpm to let proteins bind to the beads. Then, the supernatant was removed using a magnetic rack and the beads were washed off the rack 4 times with 180 µL 80% ethanol in water. Afterwards, 60 µL 50 mM ammonium bicarbonate with 1.6 µg trypsin (Trypsin Gold, Promega) and 1.6 µg LysC (mass spectrometry grade, FUJIFILM Wako chemicals) was added to the beads and put at 37°C overnight, shaking at 1000 rpm. Next day, the beads were settled on a magnetic rack and the supernatant was transferred to a new tube. The beads were rinsed with 60 µL ammonium bicarbonate and sonicated for 30 seconds, and let settle on the magnet. The supernatants were pooled and acidified with 10% TFA to a final concentration of 0.5%, then centrifugated at 20000 g for 2 min and the supernatant was transferred to a new tube. LC-MS analysis was performed on a Vanquish Neo UHPLC system (Thermo Scientific) coupled to a timsTOF HT (Bruker). The system was equipped with a CaptiveSpray ion source (Bruker), and a column oven (Sonation). Peptides were loaded onto a trap column (PepMap Neo C18, 5mm × 300 µm, 5 µm particle size, Thermo Scientific) using 0.1% TFA as mobile phase, and separated on an analytical column (Aurora Ultimate XT C18, 25 cm × 75 µm, 1.7 µm particle size, IonOpticks) applying a linear gradient starting with a mobile phase of 98% solvent A (0.1% FA) and 2% solvent B (80% acetonitrile, 0.08% FA), increasing to 35% solvent B over 60 minutes at a flow rate of 300 nl/min. The analytical column was heated to 50°C. The mass spectrometer was operated in data-independent acquisition (DIA) parallel accumulation serial fragmentation (PASEF) mode. MS2 data were acquired with eight PASEF scans per duty cycle, each containing three m/z windows. The m/z window widths were adjusted based on the expected precursor density, covering a total range of 300–1200 m/z. The ion mobility range was set to 0.64-1.42 V*s/cm, and the accumulation and ramp time was set to 100 ms. TIMS elution voltages were calibrated linearly to obtain the reduced ion mobility coefficients (1/K0) using three Agilent ESI-L Tuning Mix ions (m/z 622, 922 and 1,222). Collision energy for fragmentation was scaled linearly with precursor mobility (1/K0), ranging from 20 eV (at 1/K0 = 0.6 V*s/cm) to 59 eV at (at 1/K0 = 1.6 V*s/cm).

### Metabolomics

For metabolomics, the pellet was resuspended in ice-cold 1x PBS and transferred to a previously weighed 1.5 ml tube. The cells were again pelleted at 5’000 g for 5 minutes at 4°C and resuspended in 1 ml of ice-cold 1x PBS. This washing step was repeated once more before finally pelleting the cells and weighing the tubes again. The pellets were then finally resuspended in ice-cold MeOH:ACN:H_2_O (2:2:1, v/v) to achieve a concentration of 100 mg wet pellet weight per 250 µl solvent. The suspended cells were vortexed for 30 seconds before performing three freeze in liquid nitrogen and thaw at room temperature cycles. Proteins were then precipitated overnight at -80°C and removed by centrifugation for 15 minutes at 13,000 rpm at 4°C to generate the final supernatants for analysis. For the background sample, the same procedure was performed with a bacteria-free LB culture. Supernatants were separated on a liquid chromatography system (Ultimate 3000 HPLC system, Thermo Fisher Scientific, Germany) that was operated at a flow of 100 µL/min with an iHILIC®-(P) Classic HPLC column (HILICON, 100 x 2.1 mm; 5 µm; 200 Å; equipped with a guard column). 1 µL of sample was injected and a stepwise gradient (adapted from Wernisch and Pennathur^56^) was used for the separation procedure. The starting conditions were 90% A (100% ACN), ramp to 25% B (25 mM ammonium hydrogen carbonate, pH 8) within 6 min, 2 min hold at 25% B, ramp to 60% B from 8 to 21 min, switch to 80% B at 21.5 min, following by a flushing (21.5-26 min: 80% B) and re-equilibration step (26.1-35 min at 10% B). The HPLC was hyphenated to an Exploris 480 high-resolution mass spectrometer (Thermo Fisher Scientific, Germany) and spectra were recorded in full MS polarity switching scan mode. The ionization potential was set to +3.4/-2.8 kV and the sheath gas flow was 35. The scan range was set to 70-900 m/z and the temperature for the ion transfer tube and the vaporization were set to 300 and 250 °C, respectively. Lock mass correction in the scan-to-scan mode was applied using EASY-IC. Samples were randomly analyzed and bracketed by extraction blanks and a pooled sample QC used for background identification and instrument drift normalization, respectively. In addition, data-dependent acquisition of the QC was performed as well as the deep scan method of AcquireX was applied on the QC and selected samples to obtain additional fragmentation spectra. The raw files were analyzed with Compound Discoverer 3.3 SP3 based on an optimized untargeted workflow. The identification of the compounds was based on our in-house library obtained from authentic standard measurements. Retention time (max. 0.4 min deviation) and fragment pattern match were performed with the mass list and mzVault node, respectively. Additional fragment pattern match (at least 75) was performed with the mzCloud database from Thermo Scientific. The ChemSpider node provided information on formular and exact mass matches (max. 3 ppm) from KEGG, HMDB, E.Coli Metabolome, and BioCyc database.

### Joint Pathway Analysis

The identified metabolites and proteins with their respective fold change values in Supplementary Data 1 were uploaded to MetaboAnalyst^28^ (https://www.metaboanalyst.ca/MetaboAnalyst/upload/PathUploadView.xhtml, accessed on 30.03.2026) for Joint Pathway Analysis (JPA). A targeted metabolomics type was chosen (compound list) with the KEGG and Uniprot Protein IDs specified in Supplementary Data 1. All pathways (integrated) was chosen for the pathway database, Fisheŕs exact test was chosen for the enrichment analysis, Degree Centrality was set for the topology measure, and combine p values (pathway-level) was used as the integration method.

### Motility assay

Swimming plates consisting of 20 mL LB supplemented with 0.2% (w/v) agar were prepared the day before the actual motility assay. Strains were pre-cultured overnight in LB medium as described above before being re-diluted in fresh LB medium to an OD_600_ of 0.1. From the diluted cultures, 3 µl were pipetted into the centre of the swimming plates and plates were incubated at 37°C for 6 hours. Motility of the individual strains was assessed by measuring every hour the diameter of the visible circle of bacteria expanding from the centre.

### Isolation of cytosolic and membrane fractions

Pre-cultured strains were re-diluted in 200 ml LB medium and grown at 37°C, 165 rpm to an OD_600_ of approximately 1, when a sample was taken for western blot analysis (total, T) and the rest of the culture was cooled down on ice for approximately 3 minutes. Cells were harvested by centrifugation at 5,000 rpm for 15 minutes at 4°C. The resulting pellet was resuspended in 10 ml of ice-cold water and pelleted again at 5,000 g for 10 minutes at 4°C. The pellet was now resuspended in 3 ml of ice-cold buffer A (50 mM Imidazole/HCl pH 7, 50 mM NaCl, 1 mM EDTA, 1 mM PMSF, 250 mM sucrose) and cells were opened at 1.4 kbar in the OneShot Cell Disruptor from Constant Systems Ltd. Larger cell debris was removed from the resulting lysate by centrifugation at 5,000 g for 10 minutes at 4°C. The cleared supernatant was then further subjected to ultracentrifugation at 120,000 g for 1 hour at 4°C to separate the cytosolic fraction (supernatant) from the membrane fraction (pellet). The pellet was gently washed twice with buffer A and finally resuspended in 400 µl of the same. Aliquots of both fractions were taken for western blot analysis and mixed directly with an equal volume of 2x Laemmli buffer.

### Bacterial Two-Hybrid (BACTH) assay

The reporter strain BTH101 (F-, *cya-99*, *araD139, galE15, galK16, rpsL1 (Str r)*, *hsdR2, mcrA1, mcrB1*) for the BACTH assay was transformed with the plasmid pKNT25-ratA_Δ14_ in combination with the plasmid pUT18C carrying the individual *ubi* genes. As a positive control, the pKT25-zip plasmid was transformed in combination with the pUT18-zip plasmid. As a negative control, plasmids pKNT25 and pUT18C were employed. For each biological replicate, cultures were incubated in LB medium supplemented with 100 µg/ml ampicillin, 25 µg/ml kanamycin, and 1 mM IPTG for 24 hours at 28°C while shaking at 165 rpm. The next day, the cells of 1 ml of each culture were pelleted by centrifugation at 12,000 rpm for 2 minutes at room temperate. The resulting pellet was resuspended in 1 ml of Z-buffer (60 mM Na_2_HPO_4_, 40 mM NaH_2_PO_4_, 10 mM KCl, 1 mM MgSO_4_, 40 mM β-mercaptoethanol). Of that resuspension, a 100 µl aliquot was mixed with 900 µl 0.9% (w/v) NaCl to determine the OD_600_. To the remaining 900 µl of resuspended cells, 45 µl chloroform were added and the solution was vortexed extensively. Afterwards, the solution was incubated at 37°C for 15 minutes, followed by incubation overnight at 4°C. The next day, samples were again vortexed and incubated on ice for at least 10 minutes. Aliquots of the resulting supernatant was further diluted in Z-buffer to a final volume of 1 ml in a 2 ml tube (positive control 1:20 dilution, all others 1:5). These samples were incubated at 30°C for 5 minutes before the addition of 200 µl of ONPG (ortho-Nitrophenyl-β-galactoside, 4 mg/mL) to start the reaction. To stop the reaction, 500 µl of 1M Na_2_CO_3_ was added and the time was noted. Finally, the absorption at 420 nm was measured. For the blank solution, an aliquot of the positive control was prepared as usual with the exception that the Na_2_CO_3_ was added before the ONPG. Miller units were calculated based on the following formula:

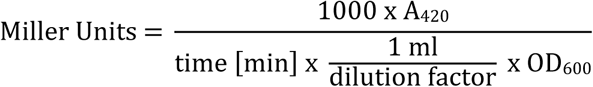

## Acknowledgments

We thank Prof. Karin Schnetz (Universität zu Köln) for gifting us the uropathogenic *E. coli* strain CFT073 and Prof. Jeff Green (University of Sheffield) for gifting us the AcnB antibody. Furthermore, we would like to acknowledge the Mass Spectrometry Facility at Max Perutz Labs for performing the proteomics analyses using the VBCF instrument pool. Both the identification of Q_8_ bound to RatA and the non-targeted metabolomic were performed by the Metabolomics Facility at Vienna BioCenter Core Facilities (VBCF), member of the Vienna BioCenter (VBC), Austria. Special thanks to Dorothea Anrather and Gerlinde Grabmann for all their help.

## Author contributions

Conceptualization, M.F. and I.M.; methodology, M.F. and I.M.; validation, M.F., L.J., D.S. and I.M.; formal analysis, M.F.; investigation, M.F., L.J., and D.S.; resources, I.M.; data curation, M.F. and I.M.; writing—original draft preparation, M.F.; writing—review and editing, L.J., D.S., and I.M.; visualization, M.F.; supervision, M.F. and I.M.; project administration, M.F. and I.M.; funding acquisition, M.F. and I.M. All authors have read and agreed to the published version of the manuscript

## Data availability

Accession numbers for the proteomics and metabolomics data respectively submitted to the PRIDE and MetaboLights repositories will be made available upon publication. All other relevant data generated in this study is included in the manuscript and supporting information. Any further requests for raw data should be addressed to the corresponding authors.

## Acknowledgment of the use of generative AI

AI was used to assist in the writing of an R script to analyze the matched features from the joint pathway analysis performed by MetaboAnalyst. During the writing process, AI was sporadically used as an interactive thesaurus.

## Supplementary Figures

**Supplementary Figure 1.**
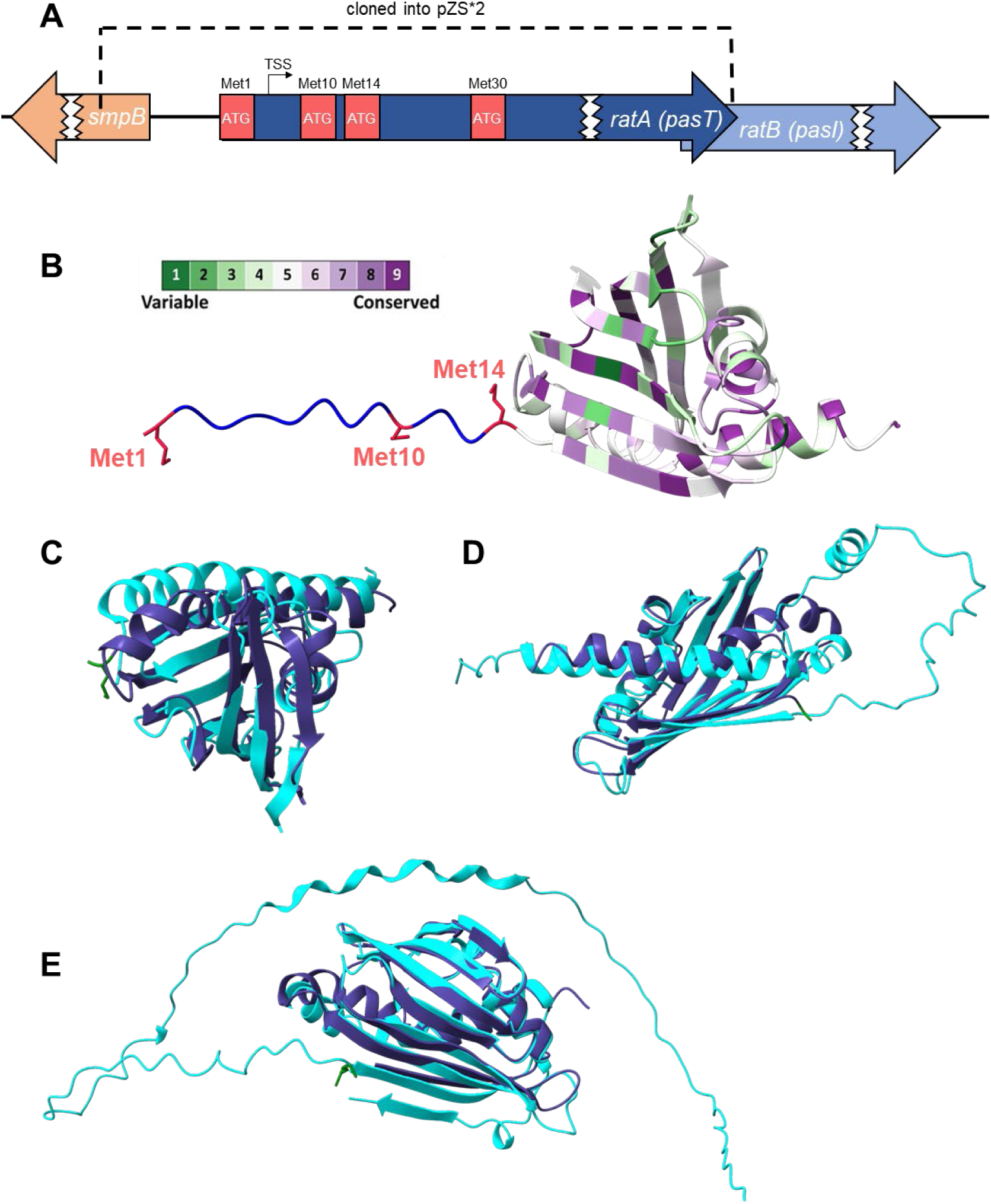
RatA structure prediction and plasmid system. **A** Not-to-scale representation of the genomic locus surrounding the *ratAB* operon as currently annotated in the K-12 reference genome (NC_000913.3). Different in-frame methionine codons at the 5’-end of the *ratA* gene are indicated in red. The transcription start site (TSS) as previously reported^23^ is indicated by a black arrow. The corresponding genomic region that was cloned into the pZS*2 plasmid for endogenous promoter control expression is marked. **B** AlphaFold^24^ structure prediction of the RatA_fl_ protein with the different methionine codons in the N-terminal extension highlighted and the conservation of the individual residues in the core START domain indicated as was calculated by ConSurf^57,58^. The same structure lacking the non-physiological N-terminal extension (dark blue) was overlaid with its homologues shown in cyan from *Caulobacter crescentus* (**C**, PDB 1T17), yeast (**D**, AlphaFold DB Q08058), and human (**E**, AlphaFold DB Q96MF6).

**Supplementary Figure 2.**
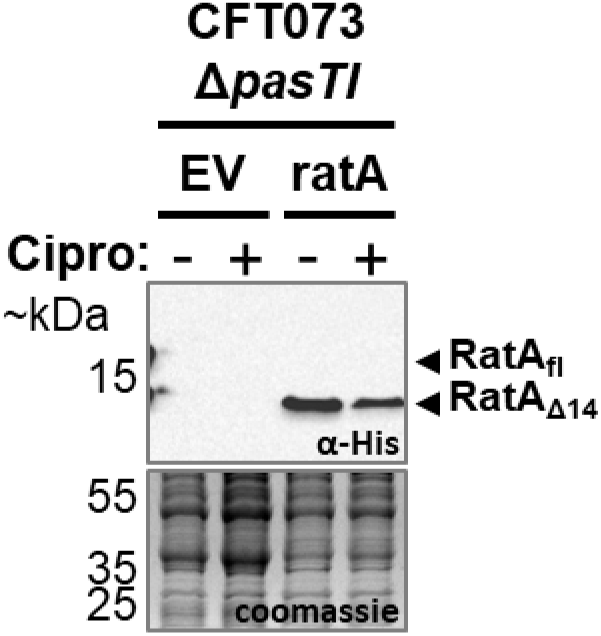
Only RatA_Δ14_ is detected in the CFT073 strain background. Western blot analysis of samples harvested from the pathogenic *E. coli* strain CFT073 with the *pasTI* operon deleted from the genome and transformed either with the pZS*2–ratA–His plasmid ectopically expressing the wildtype *ratA* gene with a 3′-terminal His-tag sequence under endogenous promoter control (ratA) or the corresponding empty vector control (EV). All strains were grown in LB medium at 37°C and antibiotic stress was induced by the addition of 10 µg/ml ciprofloxacin (Cipro) in early exponential phase.

**Supplementary Figure 3.**
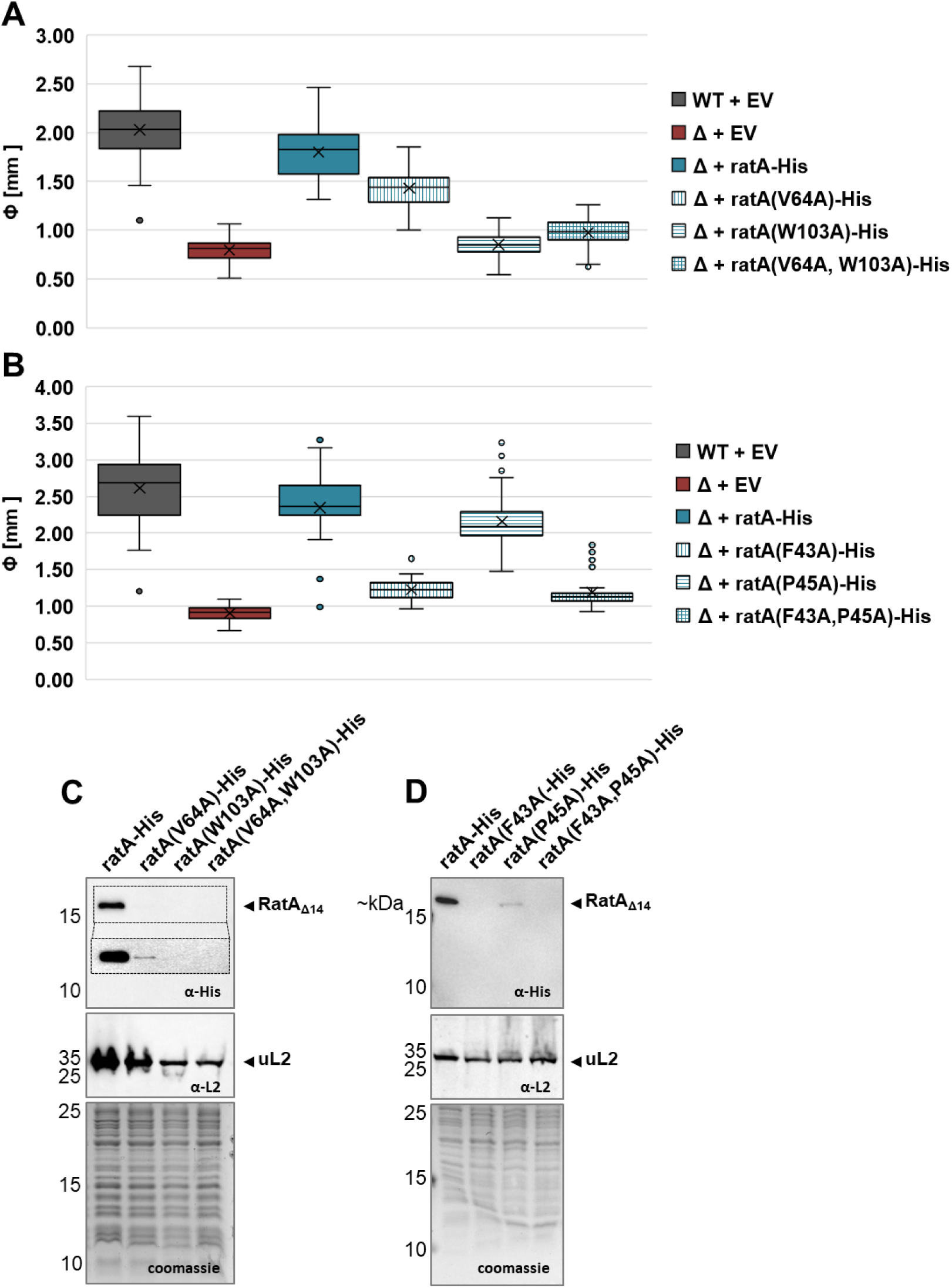
Mutations of RatA residues drastically affect the abundance of the protein. **A** Box and whisker plots representing measured colony sizes of a minimum of 30 individual colonies. Both the wildtype *E. coli* strain BW25113 (WT + EV, solid grey box) and the corresponding *ratA* deletion strain (Δ + EV, solid red box) were transformed with the empty vector pZS*2. The observed small colony phenotype was rescued when the *ratA* gene was ectopically expressed in the deletion strain under endogenous promoter control from the pZS*2-ratA-His plasmid (Δ + ratA-His, solid blue box). To investigate the influence of specific residues on the colony size, the V64A and W103A point mutations were introduced individually and simultaneously on the pZS*2-ratA-His plasmid (stripped blue boxes). **B** Same as in A, but with the introduction of the F43A and P45A mutations. **C** Protein abundances upon introduction of the same point mutations as in A were investigated by western blot analysis. The ribosomal protein uL2 served as a loading control in combination with a Coomassie stain of the whole proteome. The insert in the top panel represents the same image at a longer exposure time. **D** Same as in C, but with the same point mutations as in B and no insert for longer exposure time.

**Supplementary Figure 4.**
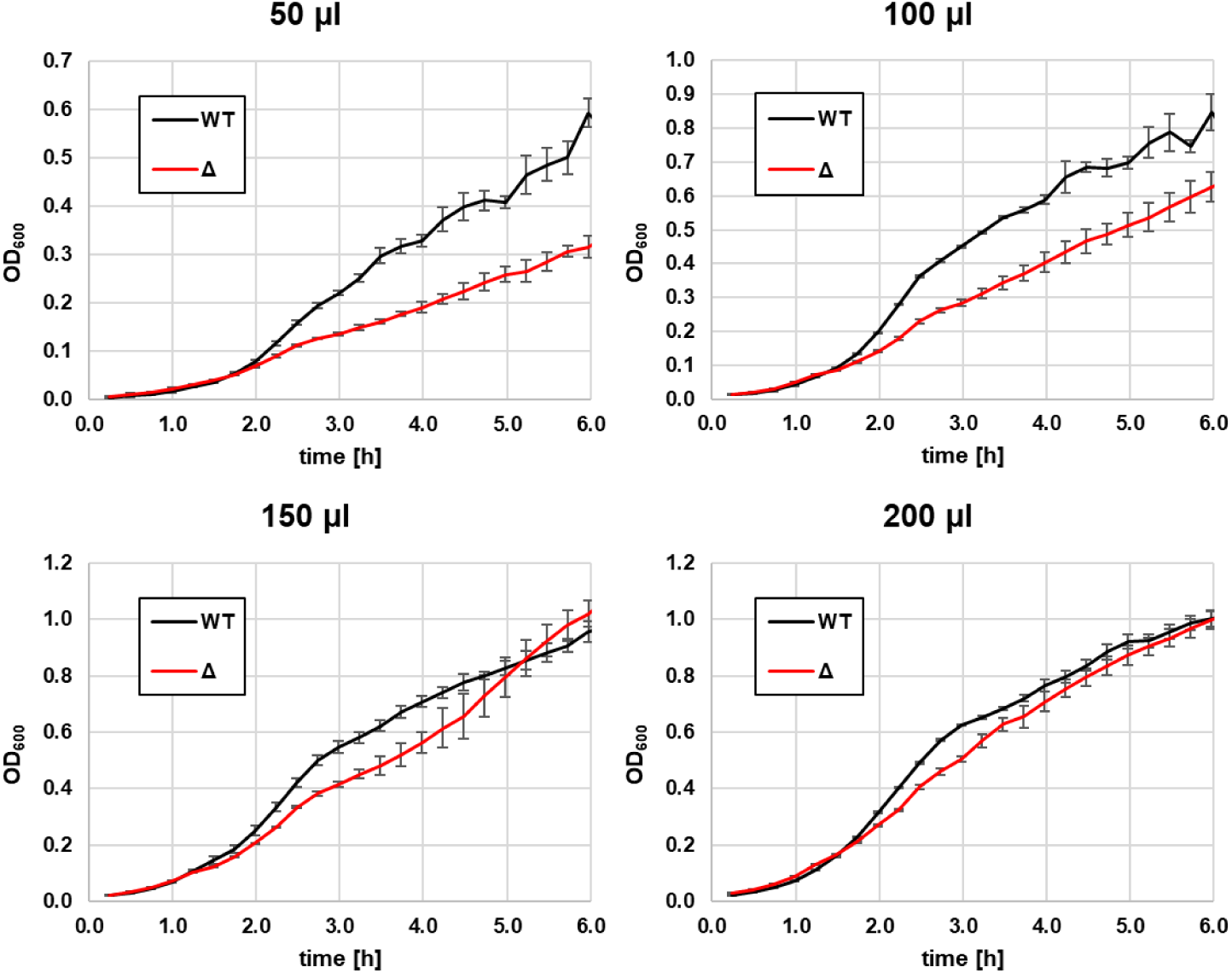
The growth phenotype depends on the volume of the culture. When grown in LB medium in a 96 well plate at 37°C while continuously shaking, the *ratA* deletion strain transformed with the empty vector pZS*2 (Δ, red line) displayed a clear growth phenotype compared to the corresponding wildtype *E. coli* strain BW25113 transformed with the same empty vector (WT, black line) only in the smaller indicated culture volumes. All panels display the average of 3 biological replicates with error bars representing the standard deviation.

**Supplementary Figure 5.**
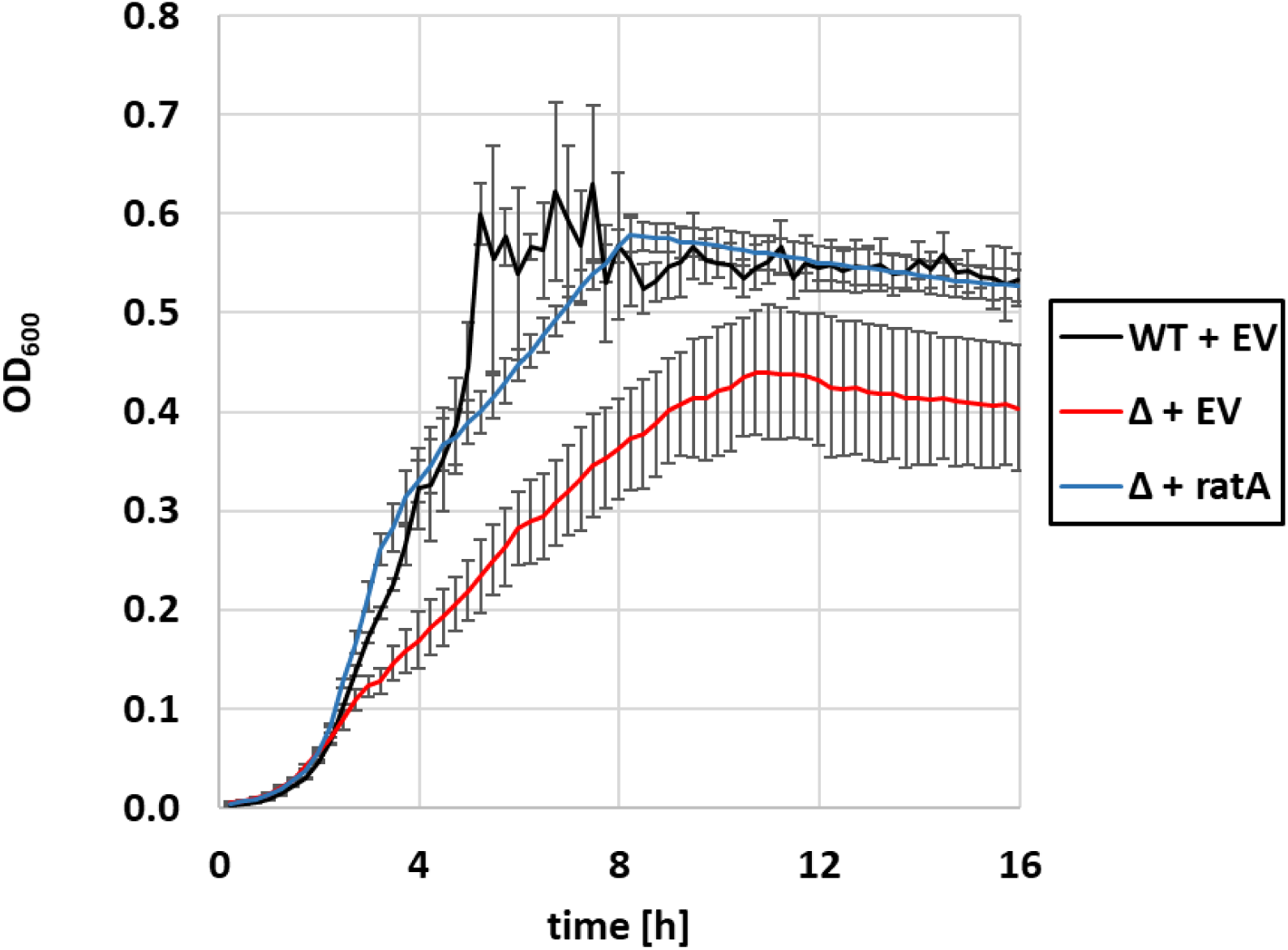
Growth of BW25113 in the presence or absence of *ratA* in a 96 well plate. When grown in LB medium in a 96 well plate at 37°C while continuously shaking, the *ratA* deletion strain transformed with the empty vector pZS*2 (Δ + EV, red line) displayed a clear growth phenotype compared to the corresponding wildtype *E. coli* strain BW25113 transformed with the same empty vector (WT + EV, black line). Ectopically expressing the *ratA* gene in the knockout strain under endogenous promoter control from the pZS*2-ratA-His plasmid (Δ + ratA, blue line) completely restored the observed phenotype. All panels display the average of 3 biological replicates with error bars representing the standard deviation.

**Supplementary Figure 6.**
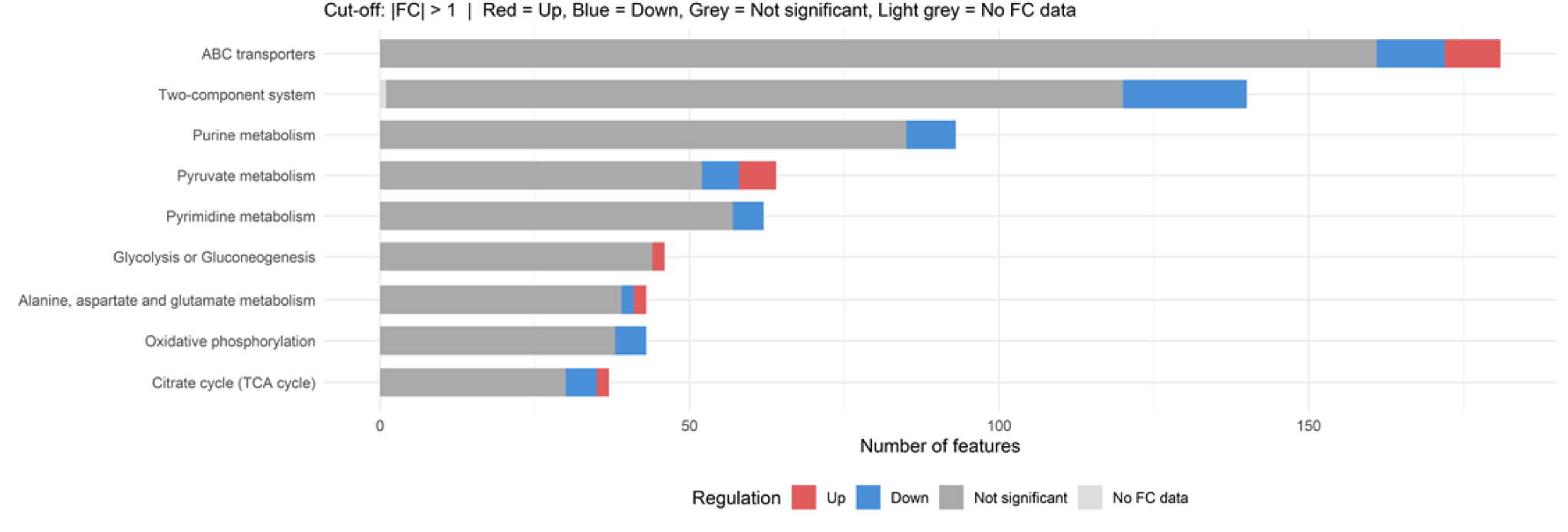
Matched features of the joint pathway analysis. Number of matched features (metabolite or gene) to the pathways highlighted in Figure 5 with coloring according to their respective up- or downregulation in the *ratAB* deletion strain (log_2_FC cut-off at +/-1). Additionally, the oxidative phosphorylation pathway was included.

